# Inflammatory and regenerative processes in bioresorbable synthetic pulmonary valves up to 2 years in sheep: Spatiotemporal insights augmented by Raman microspectroscopy

**DOI:** 10.1101/2021.04.06.438611

**Authors:** B.J. De Kort, J. Marzi, E. Brauchle, A.M. Lichauco, H.S. Bauer, A. Serrero, S. Dekker, M.A.J. Cox, F.J. Schoen, K. Schenke-Layland, C.V.C. Bouten, A.I.P.M. Smits

**Author notes:** Corresponding author: Dr.ir. A.I.P.M. (Anthal) Smits; Adress: Eindhoven University of Technology, Department of Biomedical Engineering, P.O. Box 513; 5600 MB Eindhoven.

## Abstract

*In situ* heart valve tissue engineering is an emerging approach in which resorbable, off-the-shelf available scaffolds are used to induce endogenous heart valve restoration. Such scaffolds are designed to recruit endogenous cells *in vivo*, which subsequently resorb polymer and produce and remodel new valvular tissue *in situ*. Recently, preclinical studies using electrospun supramolecular elastomeric valvular grafts have shown that this approach enables *in situ* regeneration of pulmonary valves with long-term functionality *in vivo*. However, the evolution and mechanisms of inflammation, polymer absorption and tissue regeneration are largely unknown, and adverse valve remodeling and intra- and inter-valvular variability have been reported. Therefore, the goal of the present study was to gain a mechanistic understanding of the *in vivo* regenerative processes by combining routine histology and immunohistochemistry, using a comprehensive sheep-specific antibody panel, with Raman microspectroscopy for the spatiotemporal analysis of *in situ* tissue-engineered pulmonary valves with follow-up to 24 months from a previous preclinical study in sheep. The analyses revealed a strong spatial heterogeneity in the influx of inflammatory cells, graft resorption, and foreign body giant cells. Collagen maturation occurred predominantly between 6 and 12 months after implantation, which was accompanied by a progressive switch to a more quiescent phenotype of infiltrating cells with properties of valvular interstitial cells. Variability among specimens in the extent of tissue remodeling was observed for follow-up times after 6 months. Taken together, these findings advance the understanding of key events and mechanisms in material-driven *in situ* heart valve tissue engineering.

## Introduction

Surgical or interventional valve replacement is the standard of care treatment for most patients with severe symptomatic valvular heart disease, and this treatment improves quality of life and prolongs survival. Surgical valve replacement with either mechanical or tissue valve substitutes (the latter composed of animal or human tissue and thus often called bioprostheses) generally yield favorable long-term outcomes; survival is 50-70% at 10-15 years following valve replacement [1]. Nevertheless, valve-related problems necessitate reoperation or cause death in more than half of patients with substitute valves within 10-15 years postoperatively [2], [3]. Mechanical valves induce platelet deposition and blood coagulation, (i.e., thrombosis) necessitating lifelong anticoagulation to reduce the risk of prosthetic valve-related blood clots in patients receiving them. In contrast, bioprostheses have low potential for thrombosis. However, despite improvements in tissue treatments intended to enhance durability, bioprostheses frequently suffer structural valve degeneration, often resulting from calcification, which is particularly accelerated in children and young adults [4]. Although transcatheter valve replacement technologies have recently gained favor, owing to less invasive implantation and good short-term results, their long-term durability is uncertain [5], [6]. Recently the principle of *in situ* tissue engineering (TE), also known as endogenous tissue restoration (ETR), has emerged as a promising alternative [7]–[9]. This approach utilizes the regenerative capacity of the human body to transform a resorbable polymeric implant into a living functional valve, directly in its functional site, or *in situ*. The resorbable graft functions as a suitable valve immediately upon implantation, and subsequently serves as an instructive template for progressive endogenous cell infiltration and tissue deposition [7], [10].

Preclinical studies demonstrate the potential of *in situ* heart valve TE for pulmonary and aortic valve replacements using varying materials, such as decellularized xeno- and allogenic matrix (e.g. small intestine submucosa, SIS) [11]–[13], *in vitro* cultured and decellularized matrices [14]–[17], and synthetic degradable polymers [18]–[21]. Promising functionality has been reported, as well as successful *in situ* restoration processes, including rapid cellularization, collagen deposition, endothelialization and material resorption. Recently, the first report on ongoing clinical trials using resorbable supramolecular elastomeric valves has been published, showing highly promising results when applied as pulmonary valved conduits for right ventricular outflow tract reconstruction in pediatric patients [22]. However, the regulation of the regenerative response in the complex *in vivo* environment remains poorly understood [23], and unexplained and uncontrolled adverse remodeling events such as loss of valve function, due to valve thickening and shortening, have been reported in preclinical studies (reviewed in [9]). Additionally, heterogeneity in remodeling processes, such as cell infiltration, graft resorption and ECM deposition, has been reported between valves, as well as between leaflets within the same valve [24], [25]. These variabilities in outcome emphasize the need for more in-depth knowledge of the events, kinetics and mechanisms involved in *in situ* TE, in order to achieve effective tissue formation, limit the risk of unpredicted (maladaptive) remodeling and ensure safe clinical translation.

The goal of this study was to map the long-term spatiotemporal processes of polymeric graft resorption, scaffold-induced inflammation and tissue regeneration in resorbable synthetic pulmonary valves in sheep. To that end, in depth retrospective analysis was performed on explant material of a previously reported preclinical study [19]. In that study, supramolecular elastomeric heart valve grafts (Xeltis Pulmonary Valved Conduits, XPV, Xeltis, Eindhoven, Netherlands) were implanted at the pulmonary position in an ovine model with follow-up time up to 24 months. It was demonstrated that with this graft design, safety and functionality remained acceptable throughout the follow-up time, and clinical health, blood values and systemic toxicity were not influenced by the device. Gross morphological analysis showed generally pliable leaflets, with some local anomalies, such as focal leaflet thickening or rolling of the free edge of the leaflet. The grafts were populated by endogenous cells from 2 months on, in both the conduit and the leaflet of the valves. In order to advance the understanding of how recruited cells and cellular interactions guide scaffold resorption and tissue formation *in vivo*, in the present study, grafts from the previous *in vivo* study were analyzed using a comprehensive immunohistochemistry (IHC) antibody panel [26] and Raman microspectroscopy [27][28].

The antibody panel for IHC was previously developed and validated and consists of antibodies to mark inflammatory cells, valvular interstitial cells (VICs), and extracellular matrix components, such as proteoglycans, collagens and elastic fiber-associated proteins [26]. Specifically, we assessed the presence and phenotype of inflammatory and VIC-like cells, paracrine signaling factors, endothelialization and microvascularization, and extracellular matrix components related to collagen and elastin deposition. Complementary to that, Raman microspectroscopy was applied to measure the local molecular composition of graft material and newly formed tissue in various locations of longitudinal sections of the explanted valves. Spectroscopic techniques are relatively simple, reproducible and nondestructive. Raman microspectroscopy is a vibrational spectroscopic technique that probes a specific chemical bond (or a single functional group), yielding molecular-level information of functional groups, bonding types, and molecular conformation, thus providing specific information about biochemical composition of tissue constituents and their microenvironments [29], [30]. Specifically, we applied Raman microspectroscopy on longitudinal sections of the valve explants including conduit and leaflet to assess the local chemical changes in the scaffold materials, indicative of scaffold resorption, as well as the composition and maturation of collagen in different regions of interest of the valved conduit and for various follow-up times (2, 6, 12, and 24 months). The measured trends on the molecular level were correlated to events on the cell and tissue level using IHC analysis, providing new insights into the spatiotemporal events of inflammation, tissue formation and maturation, and scaffold resorption during material-driven *in situ* heart valve tissue engineering.

## Materials and methods

### In vivo study

Explant material originates from a previously performed study (for detailed study design [19]). Briefly, a valved conduit was produced from polycaprolactone-based supramolecular polymer (electrospun polycaprolactone-ureidopyrimidinone; ePCL-UPy), material 1, and polycarbonate-based supramolecular polymer (electrospun polycarbonate-ureidopyrimidinone; ePC-UPy), material 2 (**Figure 1A**). Material 2 was designed to be more stiff when compared to material 1 to provide robustness to the conduit, while material 1 was designed to be more flexible and absorb more slowly, to accommodate the mechanical requirements as well as an anticipated slower restoration process for the leaflets. The conduits were implanted in the pulmonary position in adult Swifter sheep (age 2-4 years, weight 60-90kg) and the valves were explanted for analysis after 2 months (N=5), 6 months (N=5), 12 months (N=5), and 24 months (N=2) *in vivo*. The animal usage and study design were approved by the Test Facility’s Ethical Committee (Medanex Belgium) for compliance with regulations and the Helsinki protocol. Furthermore, animal welfare was in compliance with the Directive 2010/63/ EU.

**Figure 1:**
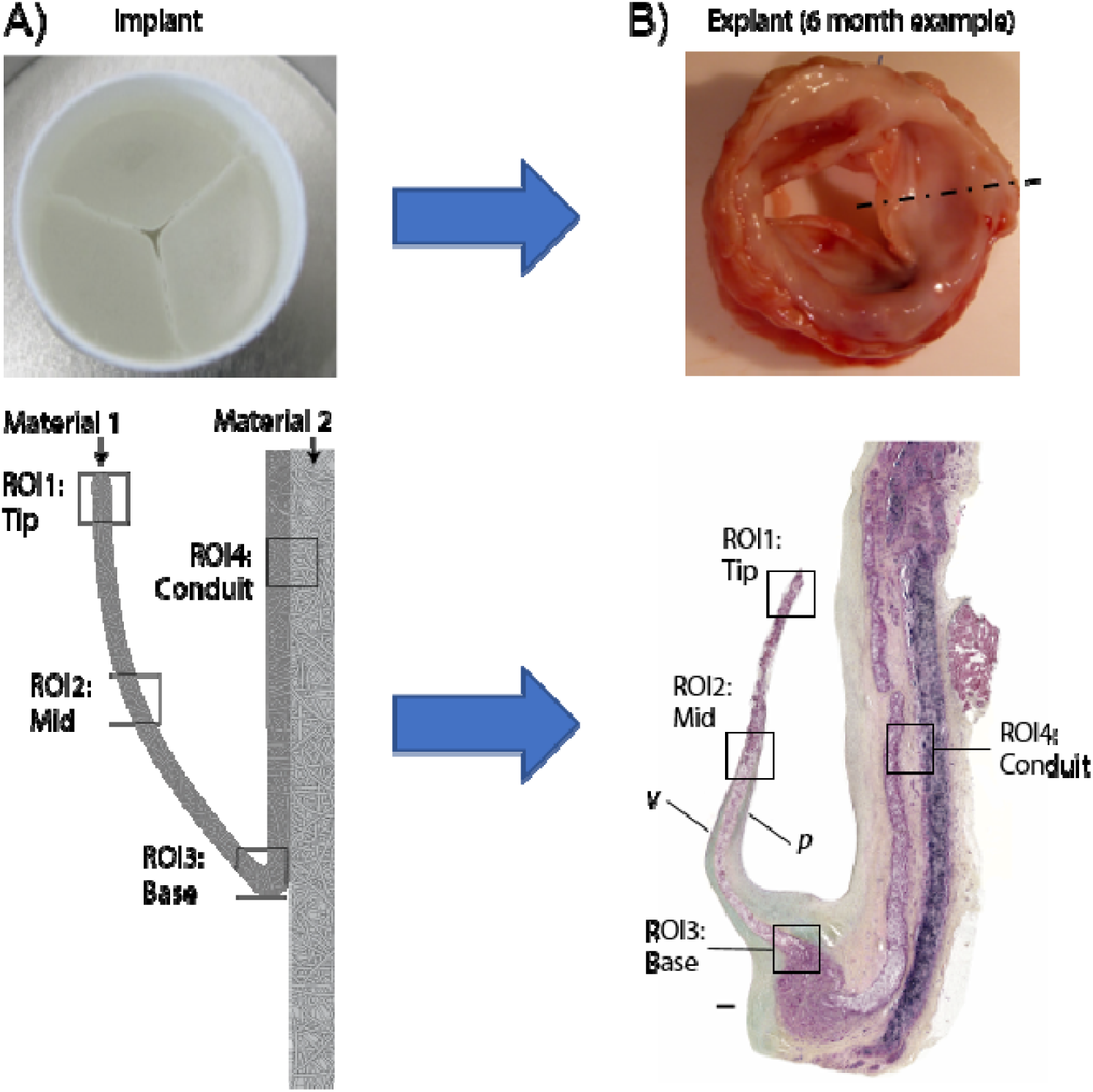
Overview of valve composition at implantation and explantation. A) Top view of the electrospun valved conduit with schematic longitudinal cross-section, indicating the composite design with material 1 (conduit + leaflet) and material 2 (conduit only), and regions of interest (ROIs) for analysis as indicated. B) Representative example of an explanted valve (6-month example) with gross morphological photograph, indicating cutting plane through the center of the leaflet, resulting in a longitudinal section through both the leaflet and conduit. Displayed is a Movat Pentachrome staining with ROIs for analysis as indicated. P and v indicate the pulmonary and ventricular surfaces of the leaflet, respectively. Photographs and Movat Pentachrome staining are adapted from[19].

### Tissue preparation

The right ventricular outflow tract including the pulmonary valve was excised and fixed in 4% neutral buffered formalin solution. The conduit was opened longitudinally and transmural specimens incorporating the leaflets and conduit wall were used for further analysis (**Figure 1B**). The specimens were dehydrated, embedded in paraffin, and sectioned at 4 to 6 microns. Sections were de-paraffinized by washing 3 times 10 minutes with xylene and rehydrated using a decreasing alcohol series. Subsequently, the sections were washed with deionized water and afterwards phosphate buffered saline (PBS, pH 7.4, Sigma).

### Immunohistochemistry

IHC was performed to evaluate the presence and phenotype of infiltrating inflammatory cells (inducible nitric oxide synthase, iNOS; CD64; CD163; CD44), VIC-like cells (α-smooth muscle actin, αSMA; embryonic form of smooth muscle myosin heavy chain, SMemb; vimentin; calponin), paracrine signaling factors (tumor necrosis factor-α, TNF-α; interleukin-10, IL-10; transforming growth factor-β_1_, TGF-β_1_), endothelial cells (von Willebrand Factor, vWF), and extracellular matrix components (collagen 1, collagen 3, (tropo)elastin, fibrillin-1, fibrillin-2) (**Supplementary Table 1**). Antibodies were selected from a previously developed sheep-specific panel [26] and performed on all explant tissues as described. The dilutions and antigen retrieval methods per antibody are specified in **Supplementary Table S1** and detailed staining protocol is specified in the **Supplementary Methods**. Briefly, biotin-labeled secondary antibodies were used and after staining, slides were either treated with SIGMA FAST™ BCIP/NBT (5-Bromo-4-chloro-3-indolulphosphate/Nitro blue tetrazolium, pH 9.5, Sigma) and counterstained with nuclear fast red (Sigma) or treated with LSAB Streptavidin and Nova Red chromogen and counterstained in Hematoxylin. For each stain, an appropriate negative and positive control tissue was selected and included, being either ovine spleen or aortic valve tissue. After dehydration and coverslipping, the stained sections were digitally scanned by CVPath at 20X magnification (CVPath institute Inc, Gaithersburg, US).

### Semi-quantitative analysis of stainings

Semi-quantitative analysis was performed for specified regions of interest (ROIs), i.e. within the graft material of the tip (ROI1), mid (ROI2) and base (ROI3) of the leaflet, the conduit (ROI4) of the pulmonary artery (**Figure 1A, B**), and the neotissue deposition onto the pulmonary artery. For each stained section, images (1.000.000 µm^2^) were selected for the ROI, as well as an image for counterstain reference and background reference using digital analysis software (QuPath) [31]. The ROIs were exported to ImageJ software (NIH, Bethesda, MD, USA) which counted pixel intensities and converted these to a histogram. Based on the intensity of the background and counterstain reference, thresholds were set in order to determine the ranges for positive staining intensity and counterstain intensity. The percentage of positive pixels was then determined by dividing the number of pixels in the positive expression range by the number of pixels in the positive and counterstain region, therefore, excluding background signal and graft material. Data were expressed as boxplots including individual data points using Prism software 5.0 (GraphPad Software, La Jolla, CA). A detailed protocol for the semi-quantitative analysis is specified in the **Supplementary methods**.

### Raman Microspectroscopy

Raman images were acquired from de-paraffinized and rehydrated tissue sections using a Raman microspectroscope (alpha300 R WITec, UIm Germany). Samples were excited using a 532 nm laser and output laser power was 60 mW. Areas of 500 x 500 µm were scanned in the following regions of interest: tip (ROI1), mid (ROI2) and base of the leaflet (ROI3) and the material interface in the conduit (ROI4) (**Figure 1A, B**). The spectral maps were collected using a line scanning mode (250 lines, 250 points/line, spectral acquisition time=0.05 s/pixel), creating a pixel size of 2 x 2 µm.

### True Component Analysis and Principal Component Analysis of Raman spectra

Raman spectra represent a highly specific fingerprint, providing molecular information on a sample [32], [33]. Multivariate data analysis tools are needed to access the comprehensive information provided by the spectral features and to generate false color-coded intensity distribution heatmaps from hyperspectral imaging maps. Here, we performed True Component Analysis (TCA) to map the location of scaffold components and biological tissue components in each sample. These components were then further analyzed for their molecular composition using Principal Component Analysis (PCA). Specifically, the scaffold material components as identified with TCA were analyzed for chemical changes indicative of scaffold resorption and the biological tissue components were analyzed for collagen formation and maturation.

Image-based data analysis was performed using Project Five Plus Software (WITec, Ulm, Germany). Of each image, data sets were pre-processed using cosmic ray removal and shape background correction. The spectra were cropped (spectral range of 250-3050 cm^-1^) and normalized to the area of 1. References spectra were measured of both non-implanted graft materials (Xeltis) and biological components, including fibrin (human plasma, sigma Aldrich #F5386), collagen 1 (lyophilized collagen 1 from porcine skin, EPC-elastin products, USA), erythrocytes, and blood components (the latter both extracted from the explant images itself). All Raman images were analyzed using TCA to identify known biological and graft material components in the recorded explant spectra based on the reference spectra (**Supplementary Figure S1**). The pixels that were recognized as similar spectra were assigned to the same spectral component and visualized as intensity distribution heatmaps to localize the different molecular components. Additionally, single point spectra of material and collagen components were extracted and analyzed using principle component analysis (PCA) using Unscrambler X14.2 (Camo, Software, Norway) to investigate molecular differences between the ROIs and between timepoints. The PCA algorithm (NIPALS) allows to decompose the spectral information deciphering spectral similarities and differences into defined vectors – so called principal components (PC). Thereby, PC-1 explains the most influencing spectral signatures, PC-2 the second most relevant information and so on. The PC scores plot can support the visualization of a separation within the dataset. The underlying spectral information is indicated in the corresponding PC loadings plot identifying bands assigned to positive score values as positive peaks and Raman shifts dominating in data with negative score values as negative loadings peaks. PC scores plots and loadings plots were employed to visualize differences of *in vivo* material and collagen spectra at different ROIs and timepoints.

## Results

### Spatial heterogeneity in inflammatory cell influx with mixed phenotypes

The *in vivo* functionality and general morphological analysis of the valved conduits up to 12 months of implantation has previously been reported by Bennink et al. [19]. The present study extends the depth of analysis of the specimens to yield a deeper understanding of the processes involved.

To augment the conventional assessment by tissue morphology of the presence and phenotype of infiltrating inflammatory cells, sections were stained by immunohistochemical methods for the general macrophage marker CD64, as well as the pro- and anti-inflammatory markers iNOS and CD163, respectively, complemented with cytokine stainings for TNF-α, IL-10 and TGF- (**Figure 2**). CD64^+^ macrophages were detected within the whole leaflet material, as well as the conduit. The distribution was heterogeneous with most abundant presence in the base region of the leaflet and in material 2 of the conduit wall, although the extend was variable between explants (**Figure 2A,B,D**). In the leaflet, less macrophages were detected in the mid region after 6 and 12 months when compared to the base and tip regions. However, an increase was seen in the mid region at 24 months (**Figure 2D**). Over time, macrophages generally remained present around and within remaining graft material, and macrophage presence diminished with full graft resorption, as was seen for example in the base region of the leaflet after 24 months. Macrophages were mainly present within and close to the graft material, however one 2 month explant was exceptional. Within the lumen of the neointima, erythrocyte-rich matrix was present in which many macrophages resided, suggesting that the influx of erythrocytes might be a cue for macrophage recruitment (**Supplementary Figure S2**).

**Figure 2:**
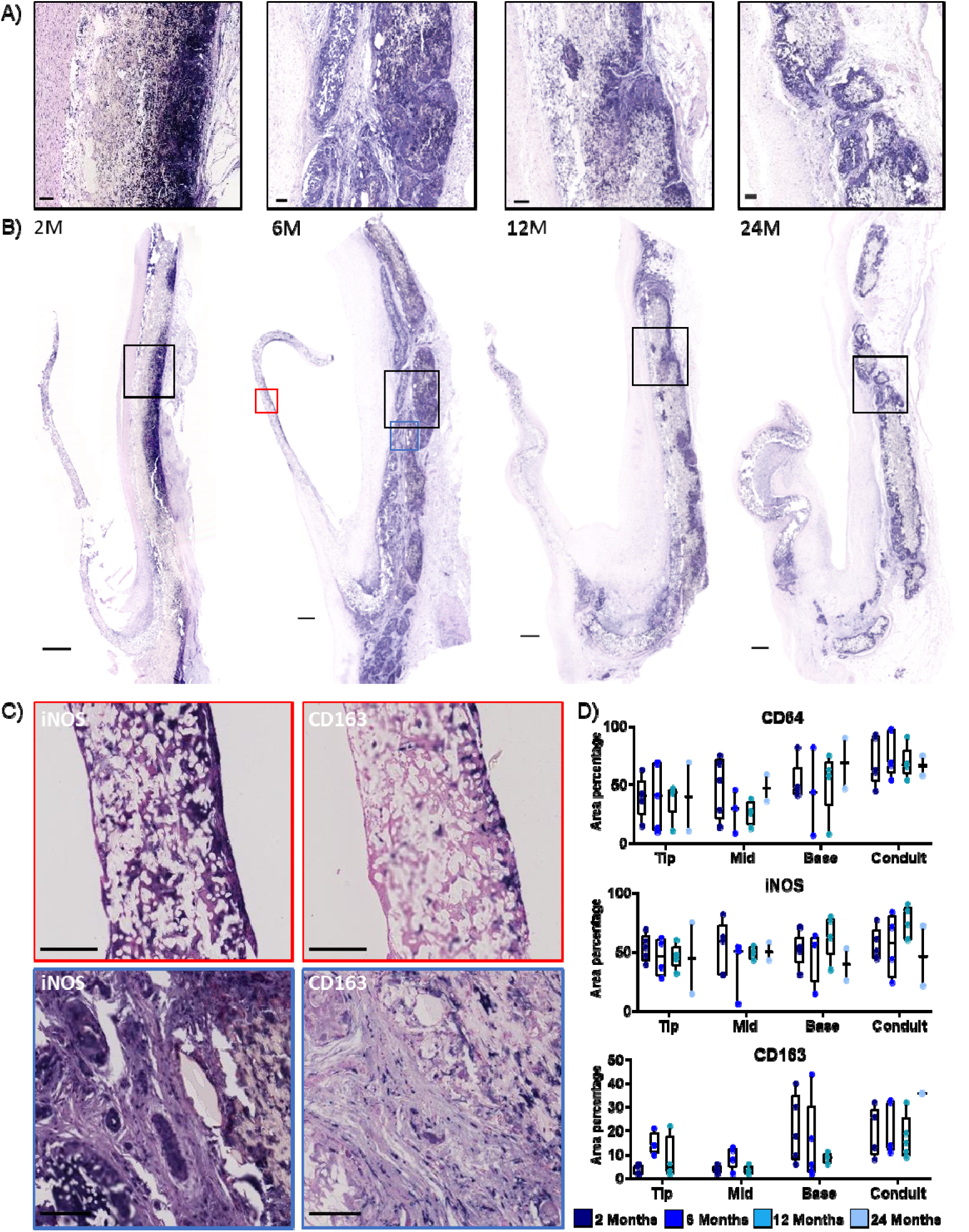
Macrophage infiltration and polarization over time is region specific. A, B) Representative images of 2, 6, 12, and 24 month explants stained for general macrophage marker CD64, with whole-valve scan and higher magnification images of areas within the graft material of the conduit (location indicated by black boxes) showing that CD64^+^ macrophages localized within both leaflet and conduit graft material. Positive staining in purple, counterstain in pink. Scale bars, 500 µm for whole-valve scan, 100 µm for zooms. C) Representative images of the pro-inflammatory marker inducible nitric oxide synthase (iNOS) and anti-inflammatory marker CD163 at the mid region of the leaflet (red boxes) and the conduit (blue boxes) of a 6 month explant showing mixed polarization with dominance of pro-inflammatory macrophages within the leaflet graft material and increased anti-inflammatory marker expression within the conduit material. Scale bar, 100 µm. D) Semi-quantification of immunohistochemical stainings (area 100.000 µm^2^). N=1 for quantification of the 24 month CD163 staining due to processing artifacts in the CD163 staining of one of the 24 month valves.

In terms of macrophage polarization and general inflammatory state, a mixed inflammatory profile of both pro-inflammatory and anti-inflammatory markers was detected throughout the implantation time within the leaflet base and the conduit (**Figure 2 C,D**). At the tip and mid regions of the leaflet, however, macrophages predominantly expressed iNOS but not CD163 at the 2 month timepoint. After 6 months, anti-inflammatory marker expression increased within the tip of the leaflet, whereas in the mid region this increase was less abundant. Expression of TNF-α and IL-10 generally correlated with the presence of iNOS^+^ pro- and CD163^+^ anti-inflammatory macrophages respectively. Additionally, TGF-β_1_ (**Supplementary Figure S3**) was mainly detected within graft material in regions with ongoing inflammation.

### Regional heterogeneity in graft resorption correlates with foreign body giant cell formation

Having established the heterogenous influx of inflammatory cells in the conduit and leaflet materials, we then analyzed the extent of graft resorption and tissue formation in the different regions of the valve. To that end, Raman microspectroscopy was performed in the specified regions of interest to acquire the molecular composition in each area. The general local composition of scaffold materials and biological components was determined via TCA. In order to analyze graft resorption per material, the spectra of material 1 and 2 were distinguished from the biological spectra using TCA (**Supplementary Figure S1**).

The spectra for material 1 and 2 were analyzed separately and compared to the pre-implant composition of each material using PCA. For the scaffold materials components, spectral changes were mainly dominated by an increasing 1123 cm^-1^ band with implantation duration in both material 1 and 2. The molecular assignment of this band is linked to C-O stretch vibrations and could be linked to an increasing contribution of alkyl-hydroxy groups referring to a degradation of the polymer. The exterior region of the conduit, consisting of material 2, demonstrated major molecular alterations as seen as an increase in the 1123cm^-1^ band already after 2 months, indicative of early resorption of material 2, with more progressive resorption over time (**Supplementary Figure S4**). The resorption of material 1 was analyzed per region of interest (**Figure 3**). Multivariate analysis of the spectra extracted from the scaffold material maps (**Figure 3A, B**) and comparison of the average score values for each animal (**Figure 3C**) demonstrated the most significant variation in the spectral signature arising between non-implanted polymer material and implanted grafts. Corresponding loadings plots depicted the most influencing spectral features (**Figure 3D**). After 2 months, spectral differences were mainly assigned to intensity differences of the overall polymer spectrum, indicating general erosion of the material rather than molecular changes of the material. Resorption of material 1 after 2 months was most pronounced in the base of the leaflet. 6, 12 and 24 months samples showed alterations in the chemical composition of the polymer with a major influence of an increasing band at 1123 cm^-1^ indicating polymer resorption, which was most pronounced in the conduit region. Most variability in chemical composition amongst explants was present after 2 and 6 months of implantation.

**Figure 3:**
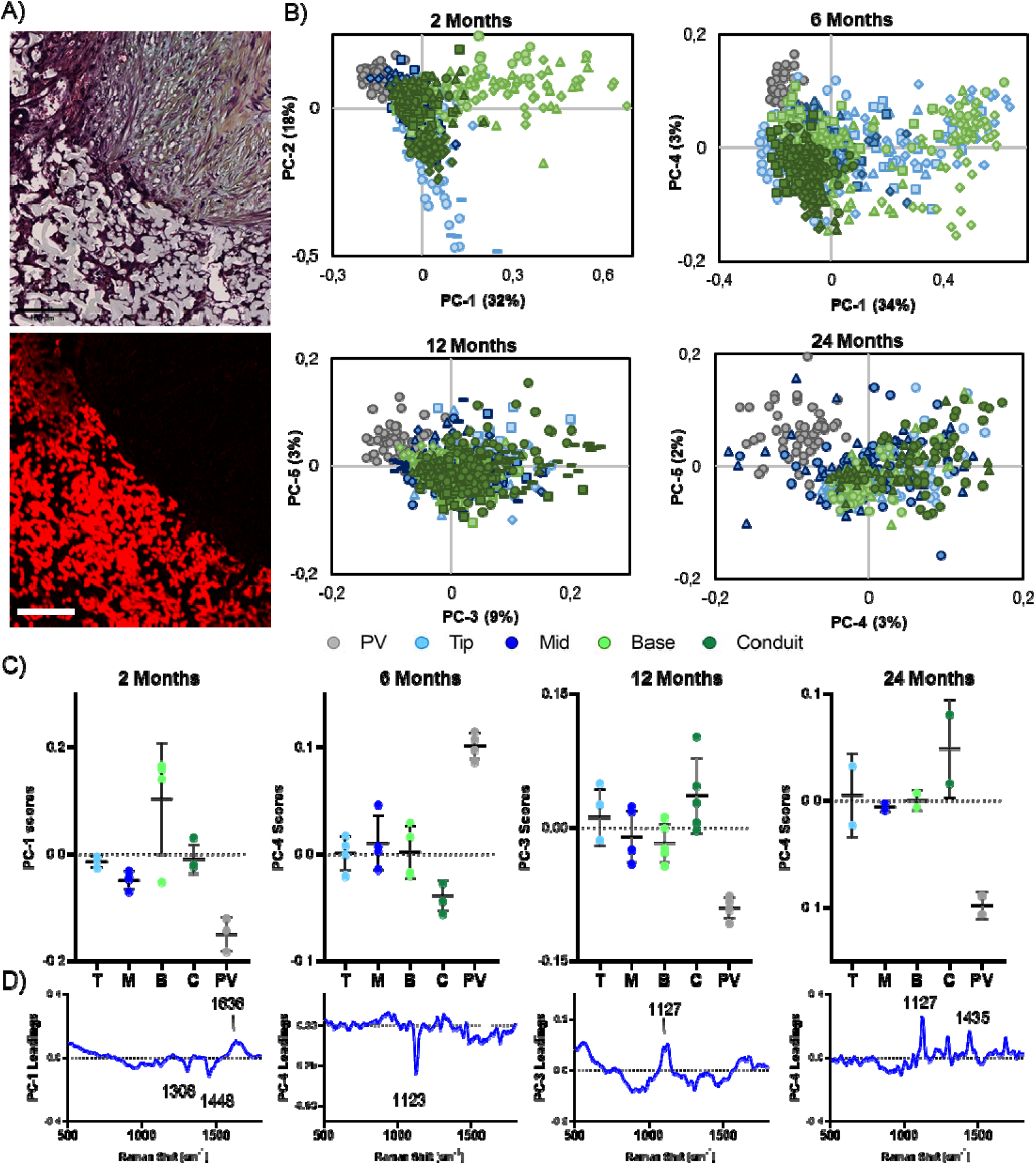
Raman analysis of *in vivo* resorption of pulmonary valve implant material 1. A) Movat Pentachrome stained base region of a 2 month explant and respective true component analysis (TCA) map of material 1 of this specific region (TCA component of material 1 depicted in red). B) Material 1 spectra were compared for different leaflet regions (ROI1-4) and non-implanted material (PV) after 2, 6, 12, and 24 months using principal component analysis (PCA) scores plot. For each PCA, spectra of the same valve are indicated with the same symbol shape. C) Comparison of ROI-specific average principal component (PC) score values (from B) for each timepoint indicated the biggest discrepancy between implanted and non-implanted material and a tendency towards the strongest resorption in the conduit region. D) Corresponding loadings plots for PCs as indicated. After 2 months, mainly differences in overall polymer signature and intensity were detected. The 6, 12 and 24 month timepoints highlighted a significant band at 1123/1127 cm^-1^ increasing upon polymer resorption.

Within the identified areas with most pronounced graft resorption, foreign body giant cells (FBGCs) were present in an inflammatory environment and remained present until full graft resorption, implying a correlation between the presence of FBGCs and graft resorption. In the conduit and leaflet base regions, FBGCs were present already after 2 months (**Figure 4A**). In the conduit, FBGCs formed predominantly at the outer boundaries of the graft material, rather than inside the graft pores, encapsulating the graft material. In the base region of the leaflet, on the other hand, FBGCs also formed within the pores of the microfibrous graft material (**Figure 4B**). Substantial variability in the overall number of FBGCs was detected in the conduit and base regions, especially at 2 and 6 months (**Figure 4A**). Few FBGCs were present in the tip and mid regions of the leaflet at all timepoints, with a slight increase over time. Heterogenous expression of both pro- and anti-inflammatory markers, iNOS and CD163 respectively, were present in FBGCs (**Figure 4C**).

**Figure 4:**
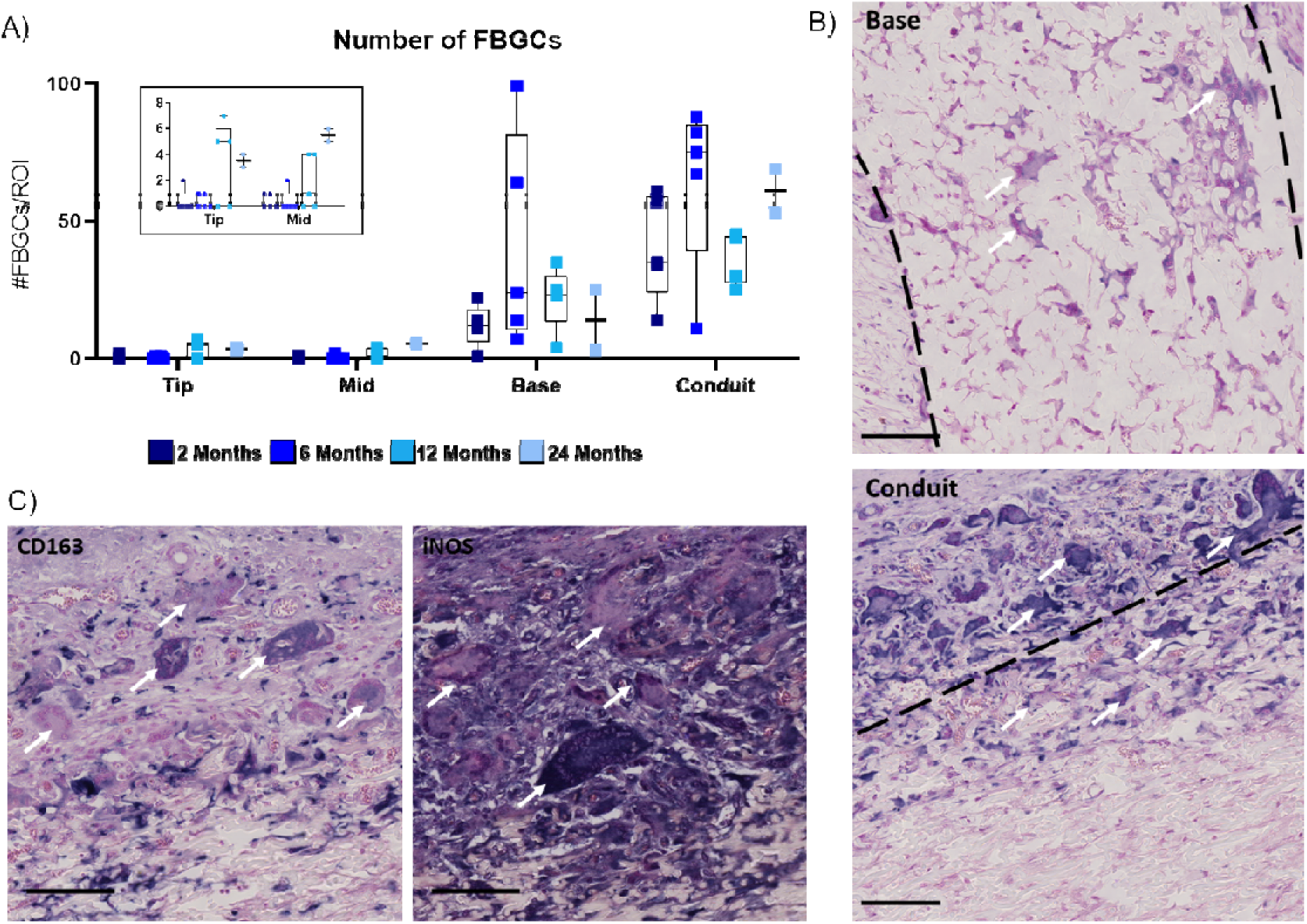
Overview of foreign body giant cell (FBGC) formation over time. A) FBGCs within each ROI (1,000,000 µm^2^) were counted in immunohistochemical images (from CD44 staining), with highest presence in the leaflet base and conduit. B) FBGCs (CD44 staining, positive staining in dark purple), indicated by arrows, within the micropores of the graft material of the base region and the outer layer of the conduit material after 2 months of implantation. Dashed lines indicating the tissue-graft interface. Scale bars, 100 µm. C) Heterogeneity between in pro- and anti-inflammatory marker expression, iNOS and CD163 respectively, in formed FBGCs, indicated by arrows, in the conduit of a 2 month explant. Positive staining in purple, counterstain in pink. Scale bars, 100 µm.

### VICs transition towards a quiescent phenotype over time

The presence and distribution of tissue producing cells was analyzed by immunohistochemical stainings for αSMA, SMemb and calponin, representing markers for (myo)fibroblast and VIC-like cell activation. At 2 months, many αSMA^+^, SMemb^+^, and calponin^+^ cells were present within the layer of neotissue covering the graft materials, as well as within the microporous graft material at the conduit and base regions specifically (**Figure 5A,D**). At this time point, SMemb^+^ and calponin^+^ cells were also present within the leaflet, whereas no αSMA^+^ cells were present in the leaflet. At 6 months, αSMA^+^ cells had migrated towards the tip of the leaflet in most explants (**Figure 5A**). At 12 months, the overall presence of VIC-like cells decreased, especially in the graft material and neotissue of the conduit and base regions. Moreover, a strong decrease in overall αSMA expression was observed between 6 and 12 months, indicative of a more quiescent VIC-like phenotype (**Figure 5D**). However, SMemb expression remained high over time. Vimentin and CD44 expression was minimal in all explant at all time points, with expression restricted to FBGCs within the graft material (**Supplementary Figure S5**).

**Figure 5:**
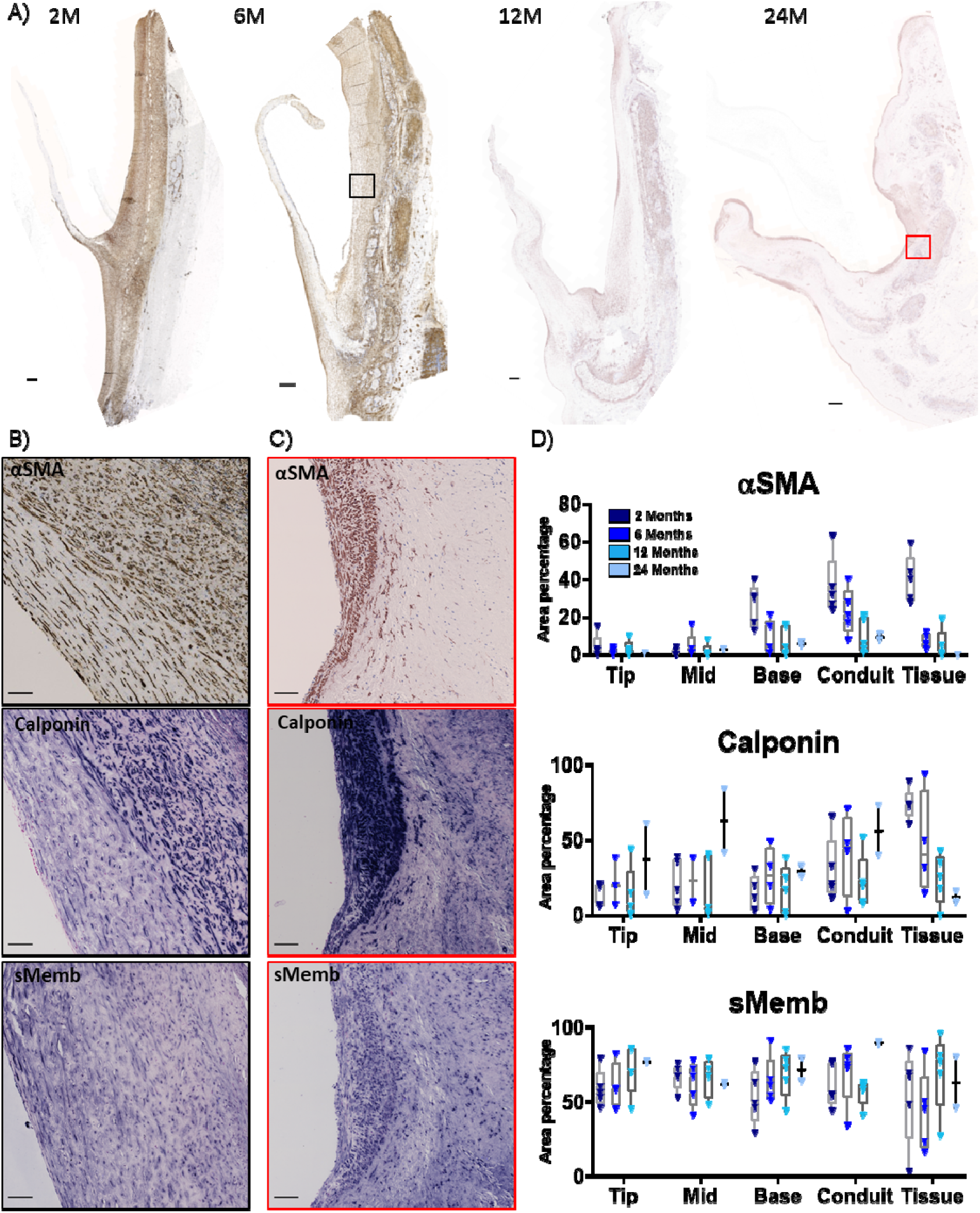
Valvular interstitial cell (VIC) phenotype shifts towards quiescent state over time. A) Overview of α-smooth muscle actin (αSMA) expression in representative explants at 2, 6, 12, and 24 months. Over time, presence of αSMA^+^ cells decreased and remaining αSMA^+^ cell localized close to endothelium. Positive staining in brown, nuclei in blue. Scale bars, 500 µm. B) and C) Heterogeneous αSMA, calponin and embryonic smooth muscle myosin heavy chain (SMemb) expression in areas with neotissue deposition onto the conduit after 6 months (B) and 24 months (C) of implantation. Locations indicated with black and red boxes in A. Calponin and Smemb; positive staining in purple counterstain in pink. Scale bars, 100 µm. D) Semi-quantification of αSMA, calponin and SMemb staining within the leaflet Tip, Mid and Base regions, in the Conduit and in the layer of neotissue deposited onto the graft material in the conduit (Tissue).

Heterogeneity in tissue deposition as well as presence and morphology of the VICs between explants and the ROIs within one explant was seen (**Figure 5B, C**). The morphology of cells varied between regions with different collagen expression, with elongated cells detected in regions that were rich in fibrillar collagen 1, and rounded cells in regions with more a dense, collagen 3-rich matrix (**Supplementary Figure S6**). Biglycan and TGF-β1 expression tended to correlate to calponin and α-SMA expression, suggesting these cells to be predominant producers of these proteins (**Supplementary Figure S6**).

### Collagen maturation between 6 and 12 months of implantation with limited elastogenesis

Having established the spatiotemporal patterns in the phenotype of tissue-producing cells, we then assessed the composition and maturation of the endogenously produced tissue. To that end, the fingerprint region of the biological component of the Raman spectra was analyzed for collagen maturation, in combination with immunohistochemical analysis. Overall, there was progressive tissue deposition in the TEHVs, in the form of pannus formation on the blood-contacting surface of the graft material as well as tissue formation in between the microporous graft fibers as seen in the TCA images (**Figure 6A, Supplementary Figure S7**). PCA analysis of the Raman spectra in the tip and mid regions of the leaflet revealed a predominant shift in PC scores between 2 and 6 months, indicating that collagen deposition within the graft material in these regions was mostly detected from 6 months on (**Figure 6A, B**). The loading plots revealed that changes over time within the leaflet tip and mid regions were dominated by increasing peaks at 860, 940 and 1250 cm^-1^ assigned to the C-C collagen backbone as well as to hydroxyproline, which are mainly linked to overall collagen content (**Figure 6A, B**). In contrast, the PC scores for the leaflet base and conduit regions showed a more gradual shift between time points, which indicates that collagen was already deposited at 2 months, followed by gradual maturation over time (**Figure 6C, D**). In the leaflet base, implantation-time dependent findings were observed, with an increasing influence of the amide I band around 1660 cm^- 1^, referring to 3D structural changes due to alterations of α-helix and β-sheet contribution which are present in more stable collagen structures [34] (**Figure 6C**). The tissue within the conduit indicated slightly different molecular alterations compared to the other regions, with collagen signatures dominated by (hydroxy-)proline at early timepoints, whereas at later timepoints signal intensities assigned to amide I (1664 cm^-1^) and III (1250 cm^-1^) as well as the C-C backbone (1450 cm^-1^) increased (**Figure 6D**).

**Figure 6:**
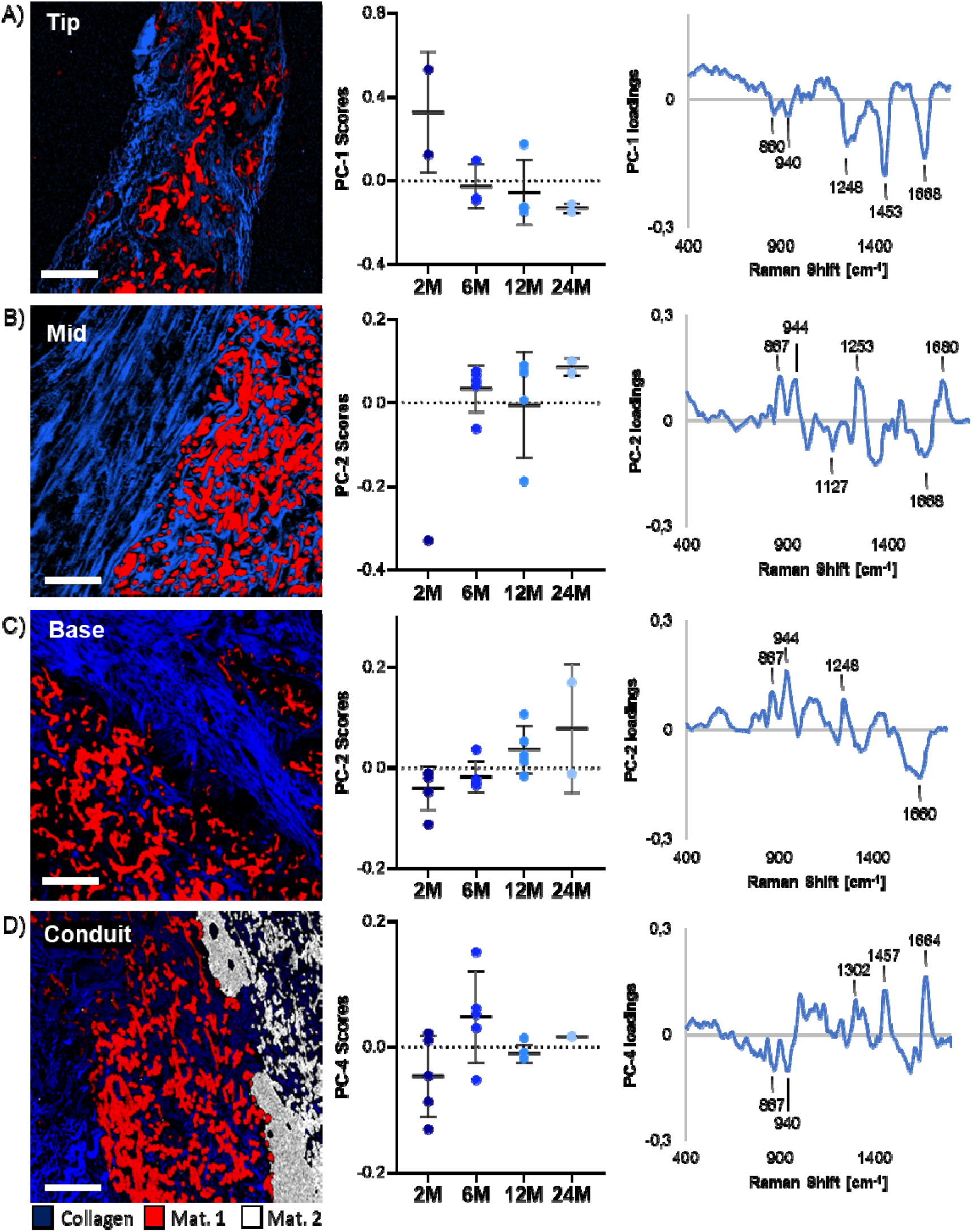
Raman microspectroscopic analysis of collagen maturation. True Component Analysis (TCA) mapping shows the localization of spectra linked to collagen (blue), material 1 (red) and material 2 (white) for the leaflet tip (A), mid (B), base (C) and conduit (D) regions. Principal Component Analysis (PCA) was performed on the collagen component from the TCA to study collagen maturation with time of implantation. Displayed are the Principal Component (PC) score plots (middle column) and the corresponding loadings plots (right column) for each ROI. Differences in leaflet tip and mid regions were mainly linked to the overall collagen content indicated by bands at 860, 940 and 1250 cm^-1^ assigned to the C-C collagen backbone and hydroxyproline. In the base and conduit additional structural changes related to shifts in the amide I band (1660 cm^-1^) were observed, indicating collagen maturation.

To evaluate the extent of collagen maturation specifically, the high wavenumber regions of Raman spectra were analyzed, as shown in **Supplemental Figure S8**. A strong shift in the PC scores was detected between 6 and 12 months in the base and mid regions of the leaflet, as well as the conduit. Peaks at 2855, 2877 and 2955 cm^-1^ characteristic for mature collagen type 1 [35] were strongest after 12 and 24 month of implantation in these regions, indicating maturation occurred predominantly between 6 and 12 months. This maturation was confirmed at the protein level by immunohistochemical stainings for collagen type 1 and 3 (**Figure 7A,B**). Specifically, the immunohistochemical analysis showed a gradual increase in collagen type 1 within the graft material over time (**Figure 7A**). In addition, a layer of collagenous neotissue was covering the graft material in the conduit region, as well as the base and mid regions of the leaflet (**Figure 7B**). Tissue deposition on the leaflet tended to occur predominantly on the ventricular side at early time points, progressing from the base region to the mid region of the leaflet. The layers of neotissue were rich in collagen 3 and low in collagen 1 after 2 months of implantation, while over time, collagen 1 expression increased. This was accompanied by the formation of more aligned, fibrous organization, indicating collagen maturation, particularly in the neotissue covering the leaflet (**Figure 7B**). Within the graft material in the base region, the collagen had a random orientation, while the neotissue at the conduit consisted of more dense collagenous tissue, when compared to the leaflet (**Figure 7B**).

**Figure 7:**
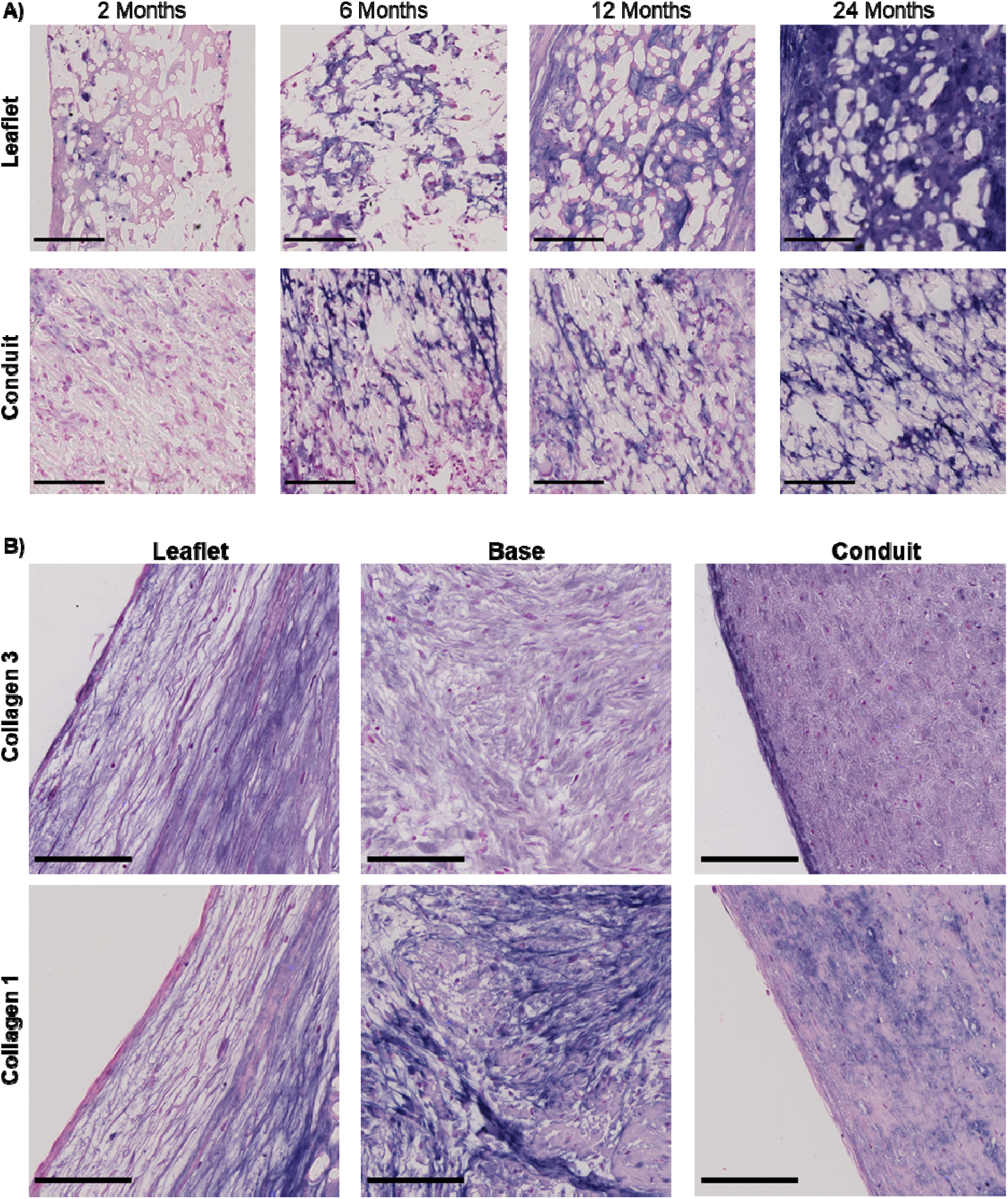
Spatiotemporal heterogeneity in collagen deposition and maturation. A) Representative images of immunohistochemical staining of Collagen 1 within graft material of the leaflet and conduit. Collagen 1 within the conduit material became more fibrillar over time compared to the leaflet. Scale bars, 100 µm. B) Representative images of collagen 3 and collagen 1, showing variations in composition and organization of collagen in the neotissue of the leaflet, base and conduit after 12 months of implantation. Scale bars, 100 µm. For all images, positive staining in dark purple, counterstain in pink.

In addition to collagen fibers, the endogenous formation of a mature elastic fiber network in the valves was assessed. Premature elastic fibers were detected at 24 months, most often located near the conduit region, as well as the leaflet base region, both at the ventricular and pulmonary side (**Figure 8A**). In all explants, (tropo)elastin was highly expressed within the layer of neotissue (**Figure 8B**). In some 6- and 12-month valves, fibrillin 2 was expressed within these regions. However, (tropo)elastin expression generally did not colocalize with the microfibrillar proteins fibrillin 1 or 2, both of which are necessary to form mature elastic fibers, as also evident from control staining of native sheep aortic valve tissue (**Figure 8C**).

**Figure 8:**
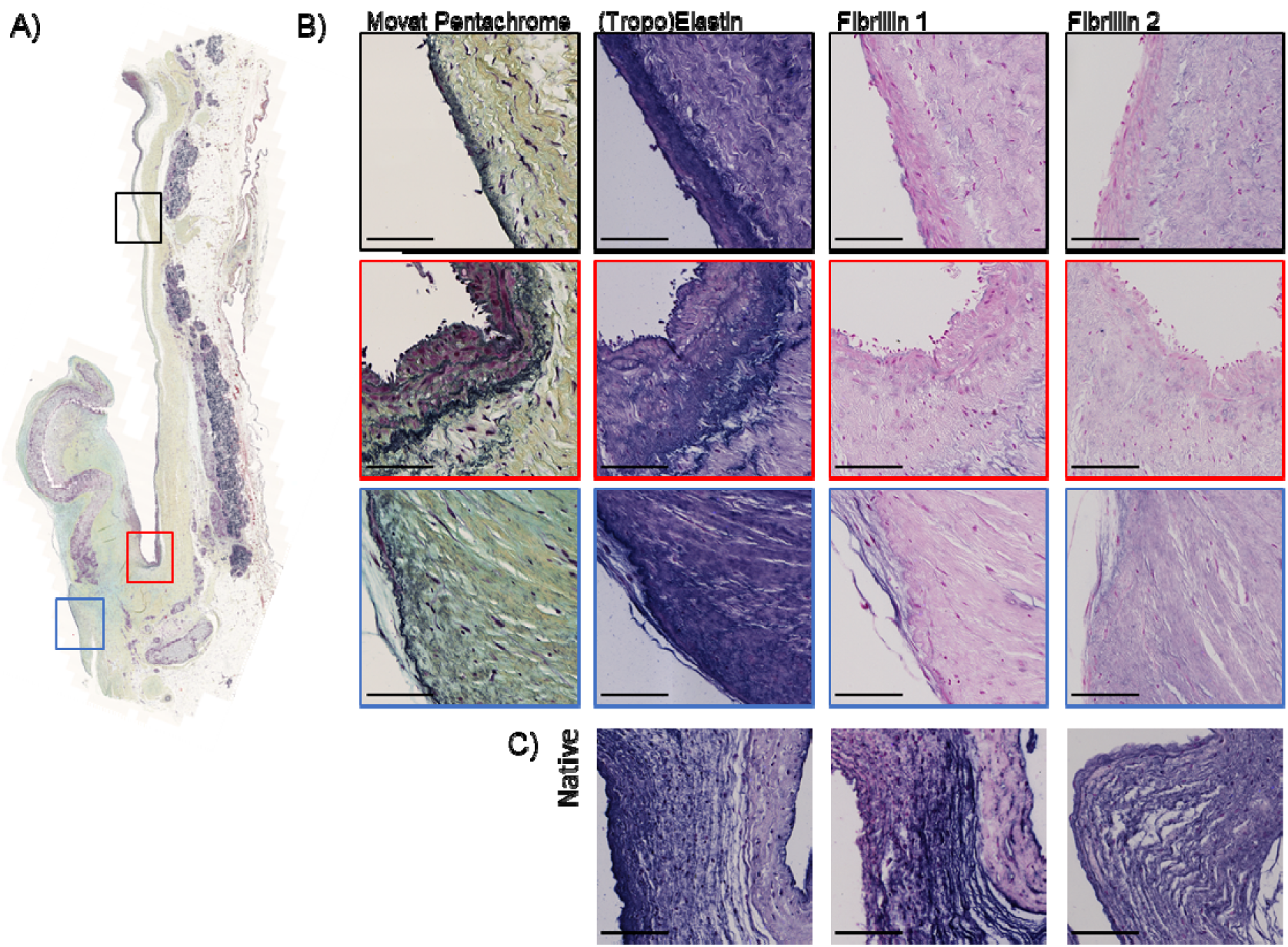
Elastic fiber network formation in 24-month explant. A) Movat pentachrome staining showing mature elastic fibers in black in a 24-month explant. Elastic fiber network formation mainly occurred near the lumen of the base region (Red and Blue zooms) and conduit (Black zoom). B) Antibody staining of the corresponding regions with (tropo)elastin, fibrillin 1 and fibrillin 2 (positive staining in purple and counterstain in pink), indicating a high expression of (tropo)elastin, but limited fibrillin 1 and fibrillin 2 expression. C) Positive control of the staining on native sheep aortic valve. Scale bars, 100 µm.

### Endothelialization and microvascularization within the microporous graft material

vWF staining was performed to study endothelialization and microvascularization. After 2 months, vWF expression was present at the luminal surface of the neotissue layer deposited on the conduit, with higher expression on the proximal side of the valve compared to the distal side (**Supplementary Figure S9**). Sparse expression was detected at the surface of the neotissue layer formed at the base region of the leaflet, predominantly at the pulmonary side and to lesser extent on the ventricular side. After 12 months, the endothelial layer was more pronounced, with expression of vWF on the neotissue of the whole leaflet, on both the pulmonary and ventricular side.

Additionally, vWF was detected within and close to the material of the conduit at all time points, indicating the formation of microvasculature within the microporous graft material (**Supplementary Figure S10**). In the leaflet, no microvessels were detected within the 2- and 6-months explants. However, the development of microvessels within the leaflet was evident in the 12 months explants, with microvessels formed at the base and mid regions of the leaflet. After 24 months, some microvessels were also present at the tip region of the leaflet. Additionally, larger blood vessels were present within the neotissue layer over the whole leaflet.

## Discussion

Within this study we aimed to gain more insight into the long-term *in vivo* inflammatory and regenerative processes in *in situ* TEHVs based on resorbable supramolecular elastomers. Enabled by a combination of Raman microspectroscopy and IHC, the main findings of this study are that: (1) inflammatory cells, and macrophages in particular, infiltrated the grafts in all regions, with a heterogenous location-dependent phenotype; (2) the resorption of the synthetic graft material was strongly location-dependent, with most rapid resorption in the conduit and leaflet base regions, and correlated with the presence of FBGCs; (3) collagen maturation was effectuated predominantly between 6 and 12 months of implantation, which coincided with a switch to a more quiescent VIC-like phenotype; (4) microvessels formed throughout the microporous graft material, forming a potential source for cell influx. Moreover, substantial valve-to-valve variability in the spatiotemporal distribution of cells, tissue and graft resorption was detected, with most pronounced variability noted in specimens following implantation of 6 and 12 months. A schematic overview of the key findings is shown in **Figure 9**.

**Figure 9:**
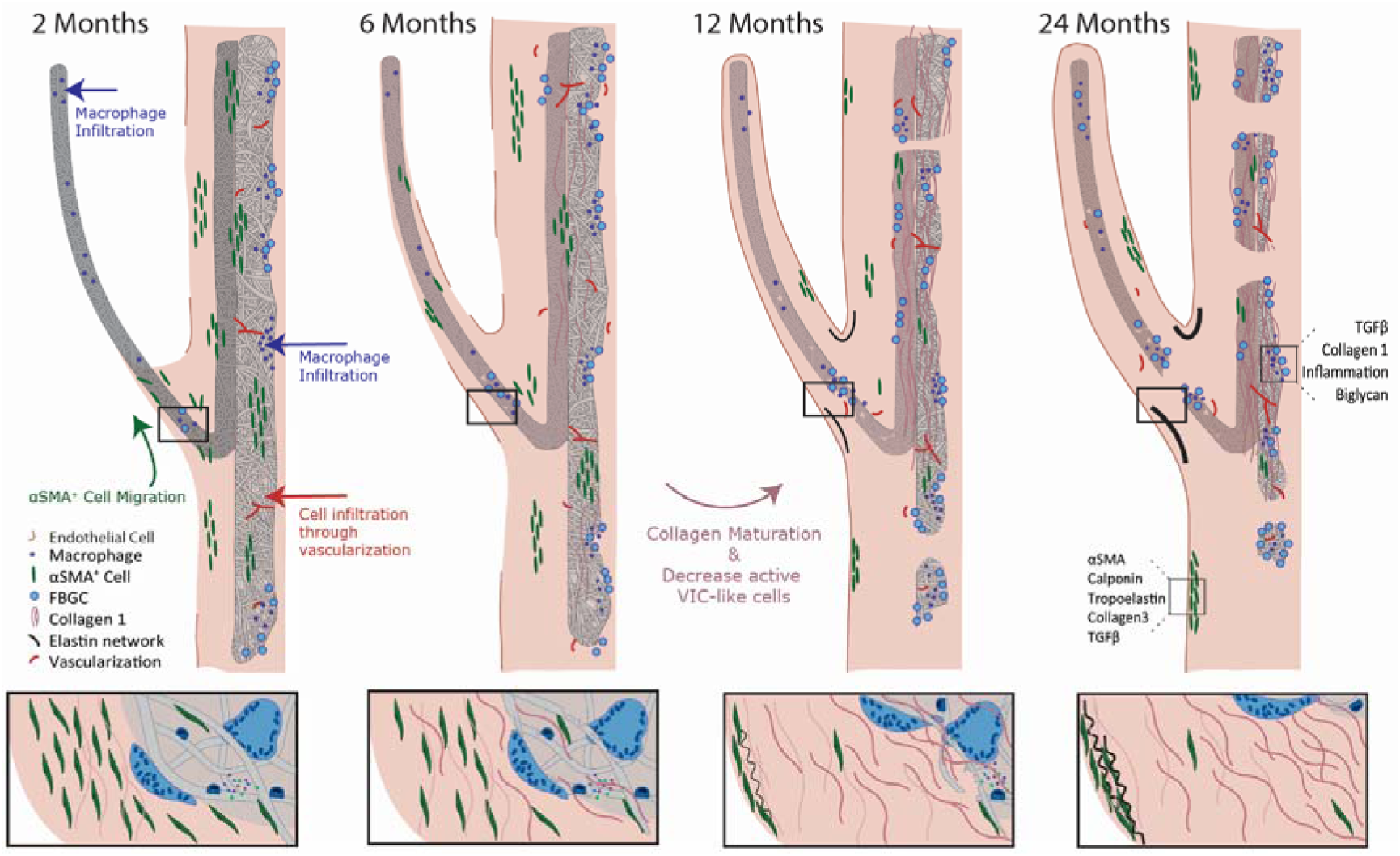
Schematic overview of essential processes involved in *in situ* heart valve regeneration. After 2 months, macrophages had infiltrated the graft material of both the leaflet and conduit. In the conduit, a microvasculature was already forming, providing an additional cell source. αSMA^+^ cells migrated from the conduit towards the tip of the leaflet over time, depositing collagen 3-rich tissue. The deposited collagenous tissue became more mature through 6-12 months, accompanied by a decrease in αSMA^+^ cells within this timeframe. Graft resorption was most pronounced in the leaflet base and conduit correlating with a high presence of FBGCs in an inflammatory environment, eventually causing successive resorption within these regions after 24 months. Inflammation had resolved in regions with complete graft resorption. Areas with ongoing inflammation were high in TGF-β_1_, collagen 1 and biglycan. Areas in which αSMA^+^ cells localized underneath the endothelium were high in calponin^+^ cells, (tropo)elastin, collagen 3 and TGF-β_1_.

Our data shed more light onto the processes of *in situ* heart valve tissue engineering. So far, the knowledge on *in situ* heart valve tissue engineering using resorbable synthetic scaffolds has been largely speculative and predominantly based on studies on *in situ* tissue-engineered vascular grafts, which have a less complex geometry and hemodynamic environment. Recent data for vascular grafts suggests that the regenerative response is driven by inflammation[36]–[38]. Macrophages, in particular, are thought to be instrumental cellular players in this process as they have a dual function: on the one hand, macrophages govern tissue formation via cross-talk to tissue-producing cells [39]–[41], while on the other hand, macrophages dictate graft resorption through oxidative and enzymatic pathways [42][43]. Here, we observed an influx of macrophages throughout all regions of the graft materials, suggesting indeed a dominant role for these cells in the regenerative cascade. Moreover, scaffold resorption was highly localized and associated with presence of FBGCs, suggesting a role for these fused macrophages in degrading the synthetic graft material through eliciting reactive oxygen species (ROS) [44]. However, the relative contributions of individual macrophages and FBGCs in graft resorption remain unclear. Interestingly, the FBGC formation in the conduit region tended to localize around the outer boundaries of the graft material, rather than within the pores of the highly porous graft material as was observed in the leaflet material. This difference may be a consequence of the differences in hemodynamics between the moving leaflet versus the static conduit wall. Alternatively, subtle differences in microstructure, e.g. fiber diameter and pore size, between leaflet material 1 and conduit material 2 may have played a role. Pore size is one of the design parameters known to influence cell infiltration, fibroblast activation as well as macrophage polarization [45]–[47]. Moreover, Wissing et al. previously showed that resorption of microfibrous scaffolds by macrophages is dependent on the scaffold microstructure [48]. *In vitro* FBGC formation is known to be regulated by multiple factors, such as cell density, material chemistry and structure, and interleukin-4 (IL-4) stimulation, with the latter being recently suggested to be most important [44], [49], [50]. In the leaflet base and conduit regions more transmural migration of macrophages is occurring and more M2 macrophages, known to secrete IL-4, are present alongside iNOS positive macrophages, when compared to the mid and tip region of the leaflet. Possibly, this contributes to a higher presence of FBGCs in the base and wall region.

Notably, the FBGC- and macrophage-rich regions were also the most active regions of cell activation and tissue formation. These activated cells had phenotypic features in common with VICs. Dependent on the mechanism of stimulation, macrophages undergo phenotypic modulation and secrete a variety of paracrine factors by which they regulate the behavior of tissue-producing cells (e.g. differentiation, migration and proliferation) and thus the deposition of new extracellular matrix [39]–[41], [51]. Importantly, it was observed that macrophages cleared from regions in which the scaffold had been completely resorbed, thereby preventing a chronic inflammatory environment that may cause excess collagenous-fibrous tissue deposition, eventually leading to valve stiffening and malfunctioning [52]. The balance between graft resorption and neotissue formation is of utmost importance in order to maintain the mechanical integrity of the valve during the regenerative response, especially within the stringent hemodynamic environment, as unbalanced remodeling could lead to early graft failure or cause prolonged inflammation and accumulation of fibrotic tissue. The Raman microspectroscopy analysis uniquely enabled the precise characterization of the molecular composition of both tissue and graft material in the same locations. Interestingly, TCA analysis suggested the presence of fibrin-like tissue, tightly surrounding the synthetic graft fibers, in both graft materials at all timepoints (**Supplementary Figure S11**). Within these regions, erythrocytes and overlap with blood components, could also be detected, both inside graft material and the neotissue surrounding the graft material. It is interesting to further define this structure, as fibrin is thought to be deposited as part of the early host response [53] and to serve as a template for new tissue formation similar to the preliminary matrix in physiological wound healing [54]. Importantly, we detected that collagen maturation developed mainly between 6 and 12 months after implantation. The immunohistochemical stainings revealed a transition from a more collagen 3 to a more collagen 1 dominated tissue, which is similar to physiological wound healing.

The formation of elastic fibers was detected only at late timepoints, with mature elastic fibers mainly detected after 24 months of implantation. In native heart valves, the elastin network is formed at the very early life stages and remains functional throughout a lifetime, with little to no new elastin formation [55]. Similar to previous studies, sparse mature elastic fiber formation was detected, although not yet comparable to the native elastic fiber network [18][56]. In all valves, (tropo)elastin was extensively produced, whereas the expression of the microfibrillar network protein fibrillin 1 and fibrillin 2 was very sparsely detected. This suggests that the bottleneck for new elastic fiber formation might not be the production of tropoelastin and the formation of elastin as such, but rather the presence and/or assembly of network proteins. We speculate that the abundant presence of inflammatory cells leads to a proteolytic environment of elastases and matrix metalloproteinases (MMPs) that may inhibit elastogenesis [57]. This may also explain why elastic fiber assembly was only detected at late time points, at which the inflammatory environment had locally resolved in graft-free regions. The latter was recently reported by Duijvelshoff et al., who demonstrated a trend of mature elastic fiber regeneration in resorbable synthetic endovascular stents in pace with a reduction in macrophage presence [58]. Overall, the present findings regarding tissue formation and maturation suggest that scaffold resorption of *in situ* TEHVs should be relatively slow in order to allow for sufficient mature and functional tissue formation prior to full scaffold resorption.

Interestingly, the maturation of the tissue was accompanied by a switch in the phenotype of the VIC-like cells to a more quiescent αSMA^-^ phenotype. In healthy adult valves mainly quiescent VICs, marked by e.g. vimentin, reside within the leaflet [59], [60], with subpopulations of smooth muscle cells and myofibroblasts [59], [61] . Surprisingly, vimentin positive cells infiltrated only very sparsely in the explants, whereas in the native sheep aortic valve used as positive control, many cells expressing vimentin were present. It is unclear what is the cause of this lack of vimentin expression. Especially at early timepoints, αSMA and calponin positive cells were abundantly present within the explants. The presence of these activated fibroblasts is typically associated with pathological remodeling and excess tissue deposition [62]. However, for *in situ* TE these cells are essential to facilitate endogenous tissue deposition. We believe it is essential that over time these cells return to their quiescent state to prevent fibrosis and leaflet contraction, as previously seen in many *in vitro* tissue-engineered heart valves [63], [64]. The marked decrease in the presence of αSMA and calponin positive cells over time, accompanied with the maturation of the tissue and the resorption of the scaffolds, indeed suggests that the regenerative response in the explants was moving towards a state of tissue homeostasis, rather than pathological remodeling. However, longer follow-up times are warranted to investigate the functionality and composition of the valves upon complete resorption of the graft materials. It also should be considered that native sheep valves have differences when compared to native human valves. Typically, ovine leaflets have a higher cellularity, more defined valvular layers and a lower amount of elastin [26], [65]. Moreover, Dekker et al. recently demonstrated that native ovine valves display a higher expression of activated VICs when compared to human valves [26]. Such differences should be considered when translating results from the ovine model towards human application.

The exact source and routes of cellular influx could not be identified from this retrospective study, and are topic of active investigation. Rapid cellularization of the leaflets of acellular valvular implants has been previously reported already 5h [66] and 15h [67] post-implantation, proposing that these cells originate from the blood. Additionally, influx of cells through transmural migration from the surrounding tissue was reported at later follow-up times [68]. Recent research suggests a dominant role of transmural migration of the surrounding tissue in small-diameter vascular graft regeneration [69]. Within the present study, after 2 months, macrophages, predominantly of a pro-inflammatory phenotype, were present within the leaflet especially near the tip, suggesting a likely origin from the circulation. Within the conduit, immune cell infiltration occurred most probably through transmural migration and neovascularization. Interestingly, calponin and SMemb positive cells were present within leaflet after 2 months, whereas αSMA^+^ cells were not found within the leaflet material but seemed to migrate from the neotissue of the wall over the leaflet material with time. The exact origin of the tissue producing cells remains unknown, however these results suggest that, similar to the infiltrating immune cells, these cells originate from various sources and from various precursor cells. In this respect, interesting phenomena were observed in the neotissue layers near the lumen, with progressively more αSMA and calponin positive cells present directly underneath the endothelium. This correlated with high levels of TGF-β and biglycan and deposition of collagen 3, which could indicate endothelium-to-mesenchymal transition (EndMT)[70], [71]. It is known that VEC-VIC interaction is very important for valve homeostasis. In adult valves, limited EndMT is taking place, and this process can contribute to replenish the VIC population as part of physiological valve remodeling [70]. However, an increased EndMT in adult valves is often associated with pathological remodeling [72]. For the analyzed explants, it remains debatable whether the process of EndMT is in fact ongoing, and if so, if it is leading to pathological remodeling or is simply part of the necessary wound healing response for example as an additional source of tissue producing cells.

One of the most important findings of this study is the spatial heterogeneity in cellular phenotypes, tissue formation and scaffold resorption. While this may partly be explained by the different sources of cell influx in the different regions, these spatial patterns also suggest an important role for the heterogeneous local hemodynamic loads. Shear stresses have been shown to play a dominant influence on the recruitment of monocytes to electrospun scaffolds [73], [74]. In addition, both macrophages and tissue-producing cells, such as (myo)fibroblasts and VICs, are known to be highly mechanosensitive [75], [76]. *In vitro* studies have demonstrated cyclic strain to influence macrophage polarization [77], [78]. Moreover, cyclic strain decreased degradative activity of macrophages within a biomaterial [78]. Additionally, both cyclic stretch and shear stress influence the cross-talk between (myo)fibroblasts and macrophages, and by that affecting tissue deposition [79]. Recent *in vivo* studies have shown the importance of hemodynamics in inflammation-driven *in situ* tissue regeneration of vascular grafts [80]–[82] and valves [16]. Moreover, in accordance with the present findings, Motta et al. recently reported on the importance of hemodynamic loads, in particular cyclic stretch, on the inflammation and *in situ* regeneration of de novo engineered extracellular matrix-based TEHVs [83]. In order to enable better prediction of *in situ* (mal)adaptive remodeling, hypothesis-driven studies have been focusing on macrophage and tissue-producing cell behavior and the influence of their microenvironment such as graft design (e.g. porosity, surface chemistry), as well as mechanical loading [16], [25], [46], [76], [79], [82], [84].

Finally, even within this very homologous test group of healthy, young sheep, variability was seen between explants of each time point. Most variability in tissue formation, as well as inflammation and graft resorption, between the valves tended to occur in the first 6 months. Within this timeframe, many processes are ongoing, influencing the timing and extend of other processes involved in endogenous tissue regenerative cascade. Potential sources of variability might be minor variabilities in the surgical procedure at implantation or through differences in host immunological state [10]. Reasonably, more variation can be expected when translating towards more heterogenous patient populations. Additional sources of variability are proposed to include genetic variation, co-morbidities and sex-and age-related changes, medications and environmental factors [10][85].

## Limitations

This study has several limitations. One important limitation is that this was a retrospective study and only endpoint data measurements were used in this study. Therefore, it remains speculative which early events could have led to certain downstream remodeling. Spectral analysis of Raman data was performed only within the graft material. As longitudinal analysis of a single valve was not possible, variability between selected areas could occur due to resorption of graft material, especially in the base region. Therefore, resorption in this region seemed less pronounced when comparing Raman analysis and immunohistochemical visualization. Additionally, intense signal and close material interactions, such as fibrin, might cause noise in the spectrum. A more detailed pixel size could be used to overcome this limitation, however, here we opted for compromising on the minimal pixel size in order to enable scanning of various larger regions within the valve. Another limitation of this study is the limited sample size. However, there are only few studies on long-term *in vivo* implantations of resorbable synthetic heart valves available to date, so despite the limited sample numbers, the presented information represents a valuable contribution to our understanding of *in situ* heart valve tissue engineering.

## Conclusion

With these analyses, we uncovered new insights in the endogenous tissue restoration response regarding infiltration and presence of immune cells and tissue forming cells and their correlation to graft resorption and neotissue formation and maturation over time. Our findings show important spatial heterogeneities in the regenerative process, most likely as the combined effect of heterogeneous cellular repopulation mechanisms and heterogeneity in the local hemodynamic loads, which may fuel the rational design of improved grafts. To fully understand the *in situ* regenerative response, more in-depth, comprehensive *in vivo* analysis and mechanistic *in vitro* work with focus on the role of local hemodynamics and the downstream effects thereof are warranted. However, to ensure robust clinical translation, a holistic approach is needed, focusing not only on the graft properties, but also on implant-independent, patient-specific factors.

## CRediT author statement

**Bente de Kort**: Investigation, Formal Analysis, Data curation, Visualization, Writing–Original Draft; **Julia Marzi**: Investigation, Formal analysis, Visualization, Writing-Review & Editing; **Eva Brauchle**: Investigation, Formal analysis; **Arturo Lichauco**: Software, Validation; **Hannah Bauer**: Validation, Writing-Review & Editing; **Aurelie Serrero**: Resources, Writing-Review & Editing; **Sylvia Dekker**: Resources, Validation; **Martijn Cox**: Conceptualization, Resources, Writing-Review & Editing; **Fred Schoen**: Validation, Writing-Review & Editing; **Katja Schenke-Layland**: Resources, Writing-Review & Editing; **Carlijn Bouten**: Writing-Review & Editing, Funding acquisition; **Anthal Smits**: Conceptualization, Formal Analysis, Writing–Original Draft & Editing, Funding acquisition, Supervision.

