## Supplement for "Inflammatory and regenerative processes in bioresorbable synthetic pulmonary valves up to 2 years in sheep: Spatiotemporal insights augmented by Raman microspectroscopy"

a. Department of Biomedical Engineering, Eindhoven University of Technology, Eindhoven, The Netherlands
b. Institute for Complex Molecular Systems (ICMS), Eindhoven University of Technology, Eindhoven, The Netherlands
c. Department of Women's Health, Research Institute of Women's Health, Eberhard Karls University Tübingen, Tübingen, Germany
d. NMI Natural and Medical Sciences Institute at the University of Tübingen, Reutlingen, Germany
e. Cluster of Excellence iFIT (EXC 2180) "Image-Guided and Functionally Instructed Tumor Therapies", Eberhard Karls University Tübingen, Tübingen, Germany
f. Xeltis B.V., Eindhoven, The Netherlands
g. Department of Pathology, Brigham and Women's Hospital, Harvard Medical School, Boston, MA, USA
h. Department of Medicine/Cardiology, Cardiovascular Research Laboratories, David Geffen School of Medicine at UCLA, Los Angeles, CA, USA

### Extended Methods

#### Immunohistochemistry (IHC)

Sections were de-paraffinized by washing 3 times 10 minutes with xylene and rehydrated using a decreasing alcohol series. Subsequently, the sections were washed with deionized water and afterwards phosphate buffered saline (PBS, pH 7.4, Sigma). After rehydration, the selected antibody retrieval method (**Supplementary Table S1**) was performed. For enzymatic antigen retrieval, 0.05% pepsin (Sigma) in HCl (8mM) was added on tissue sections for 10 min. at 37 °C. For heat-mediated antigen retrieval, tissue sections were incubated in modified citrate buffer (pH 6.1, DAKO) or TRIS-EDTA buffer (pH 9.0, DAKO) at 96 °C for 20 min. Afterwards the sections were allowed to cool to room temperature for 45 min. Non-specific antigen binding was blocked using 5% goat serum (Invitrogen) or 5% horse serum (Life Technologies) in PBS/0.05% Tween-20 (Merck) with 1% bovine serum albumin (BSA, Sigma) for 1h at room temperature. The primary antibody was added in the desired concentration (**Supplementary Table S1**) in 1:10 diluted blocking solution and incubated overnight at 4 °C. The negative control was incubated with 1:10 diluted blocking solution. The sections were washed with PBS/Tween-20. The biotin-labeled secondary antibodies (goat-anti-rabbit (DAKO) or horse-anti-mouse (DAKO)) were diluted 1:500 in PBS/Tween-20 and incubated for 1 h at room temperature. Sections were washed with TRIS-buffered saline (TBS)/0.05% Tween-20 and incubated with the ABC-alkaline phosphatase kit (Vector laboratories, VECTASTAIN ABC-AP Staining Kit) for 1h at room temperature to enhance staining. Sections were washed with TBS/Tween-20 and treated with SIGMA FAST™ BCIP/NBT (5-Bromo-4-chloro-3-indolulphosphate/Nitro blue tetrazolium, pH 9.5, Sigma) for 5 min up to 1h depending on staining intensity, considering positive and negative controls. Nuclei were counterstained using nuclear fast red (Sigma) for 5 min. After dehydration, the sections were mounted in Entellan (Merck).

For staining of α-SMA an vWF the deparaffinize sections were transferred to distilled water and EDTA antigen retrieval was performed. Endogenous enzyme activity was blocked with 3% $H_2O_2$ and Avidin/Biotin. Fc binding and charged sites were blocked with 1% BSA in PSA. Incubation with primary antibodies (rabbit antibody for vWF and mouse antibody for α-SMA) and secondary antibodies (Goat anti-Rabbit and Horse anti-mouse) was performed, respectively. Slides were incubated in LSAB Streptavidin and developed in Nova Red chromogen. Hematoxylin was used as counterstain. After dehydration, the slides were coverslipped.

For Movat Pentachrome staining the deparaffinize sections were transferred to distilled water, rinsed in 1% Acetic Acid and stained with Alcian Blue. Afterwards the slides were washed in tap water and placed in Working Elastin Hematoxylin Solution. The slides were rinsed in tap water and differentiated in 2% aqueous Ferric Chloride. After again rinsing in tap water, the slides were placed in 5% Sodium Thiosulfate and then in tap water. Now the slides were stained with Woodstain Scarlet, rinsed in 0.5% Acetic Acid and placed in 5% Phosphotungstic Acid. After rinsing in 0.5% Acetic Acid, the slides were changed of absolute alcohol and stained in Alcoholic Saffron Solution. After dehydrated, the slides were coverslipped.

**Semi-quantitative analysis of IHC stainings**

Regions of interest (ROIs) were created as square annotations of $1,000,000^2$ um in QUPath. The leaflet tip ROI was designated first as the functional tip of the leaflet, disregarding any curling or retraction of the original scaffold tip within the tissue of the leaflet tip (**Extended Methods Figure 1A,C**). The annotation was placed to maximize the amount of the functional leaflet tip present in the annotation. The leaflet base ROI was designated as the area of the leaflet marked by the observed hinge of the tissue of the leaflet and the conduit. The ROI was placed on along the middle of the leaflet base to include scaffold material and tissue along both the pulmonary and ventricular side of the leaflet. The distance between the leaflet tip and base was then measured, and the leaflet middle ROI annotation was placed in the middle of this measurement.

A semi-quantitative analysis was then performed on the leaflet ROIs by determining the ratio of pixels showing expression compared to all of the biological tissue present in a given leaflet ROI. Leaflet ROIs were then exported to ImageJ, which was then used to average the RGB values of each pixel to convert them to grayscale. Pixels in the grayscaled image were then plotted onto a histogram according to their intensities (**Extended methods Figure 1B**). As the exported ROIs were 8-bit images, the number of possible intensities is calculated as $2^8$ which allows for an intensity range of 0-255, with 0 being the darkest pixels, and 255 being brightest. For each sample, an annotation in which only pixels which were visibly counterstain was selected. These counterstain annotations were not square shaped, but were also at least $1,000,000^2$ um in total area. When the pixels within these annotations were grayscaled and plotted on an intensity histogram, they provided an expected intensity range for pixels showing only the counterstain color agent. This allowed for two thresholds to be created at the edges of this range for that particular image. Any pixels present in a given ROI within that image which were darker than the lower threshold of the counterstain range are counted as showing expression. Any pixels that are brighter than the higher edge of the counterstain range are counted as either background or fibers. The total number of pixels darker than the expected counterstain range are then divided by the total number of pixels, excluding any pixels counted as background or fibers. This exclusion of these high intensity pixels eliminates pixels of the image background as well as intact scaffold fibers, which are unstained by the IHC protocols, thereby avoiding potential reduction of perceived expression of biological material within intact fibers when evaluated subjectively by eye. It should be noted this method calculates how widespread expression is within an ROI (surface coverage), not how strong the expression (intensity of staining) in the ROI may be. For a given pixel showing expression, the intensity of that pixel does not matter as long as it is lower than the threshold set by the counterstain range. 100 pixels with an intensity value of 0 are counted the same as 100 pixels with an intensity value of 1.

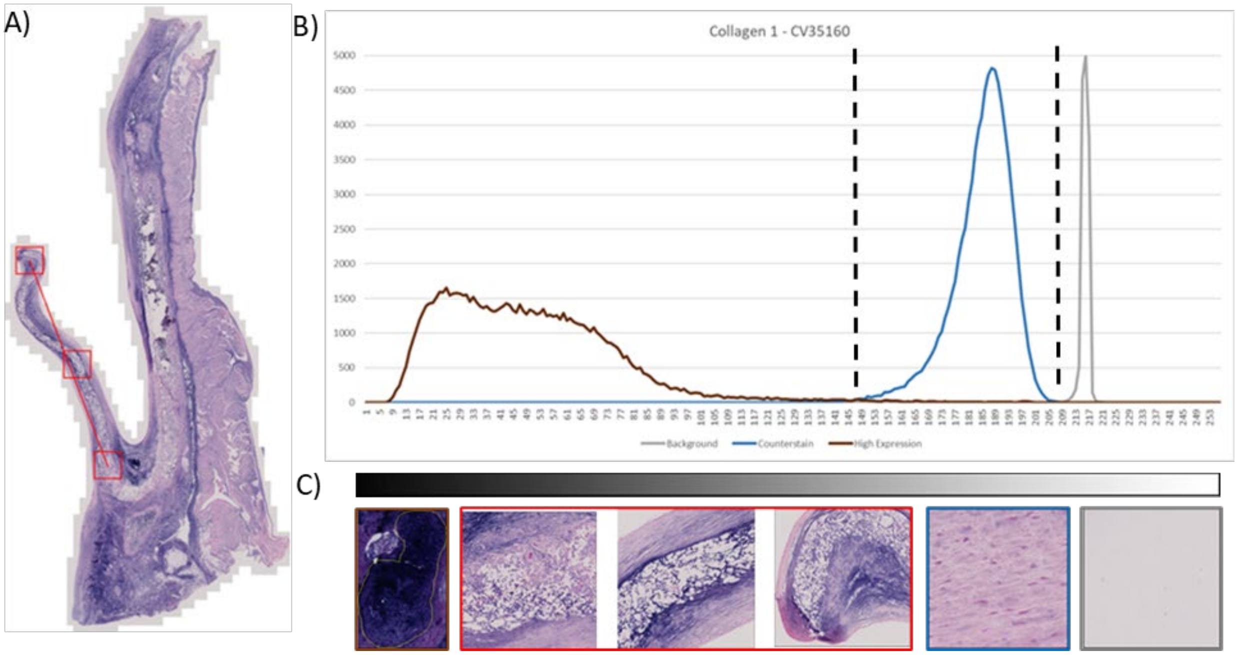


**Extended Methods Figure 1**: A) example of a 12-month explant stained for Collagen 1 as used for semi-quantification. Within the image 3 ROIs were selected (red boxes) and additionally sections with high expression, counterstain and background signal were selected. B) Intensity histogram of selected sections of high expression (brown), counterstain (blue) and background signal (grey). Dashed lines indicating thresholds for excluded backgrounds signal (right) and counterstain (left). C) Images used for semi-quantification of ROIs (Red) and to define high expression (brown), counterstain (blue) and background signal (grey).

#### Supplementary Table

**Supplementary Table S1.** Overview of selected IHC antibodies and their respective antigen retrieval methods and dilutions.

| Antibody | Description | Host | Clone | Distrubutor | Cat. No. | Antigen retrieval | Dilution |
|---|---|---|---|---|---|---|---|
| | ECM | | | | | | |
| **Collagen 1** | | Mouse | col-1, IgG1 | Sigma | C2456 | Citrate | 200 |
| **Collagen 3** | | Rabbit | Polyclonal | Abcam | Ab7778 | Pepsin | 250 |
| **(Tropo)Elastin** | | Rabbit | Polyclonal | Abcam | ab21610 | Pepsin | 400 |
| **Fibrillin-1** | | Rabbit | Polyclonal | Sigma | HPA017759 | Pepsin | 100 |
| **Fibrillin-2** | | Rabbit | Polyclonal | Sigma | HPA012853 | Pepsin | 100 |
| **Biglycan** | | Rabbit | Polyclonal | Biorbyt | Orb100396 | Pepsin | 150 |
| | Tissue cells | | | | | | |
| **Vimentin** | Fibroblasts | Rabbit | D21H3, IgG | Cell signaling | 5741 | Citrate | 400 |
| **Calponin** | Myofibroblasts/ smooth muscle cells | Rabbit | Polyclonal | Abcam | ab46794 | Pepsin | 150 |
| **SMemb** | Myofibroblasts/ Dedifferentiated smooth muscle cells | Mouse | 3H2, IgG2b | Abcam | ab684 | Tris/EDTA | 200 |
| **α-SMA** | Myofibroblasts/ activated VICs | Conducted by CVPath | | | | | |
| **vWF** | Endothelial cells | Conducted by CVPath | | | | | |
| | Immune response | | | | | | |
| **CD64** | Pan-macrophage | Mouse | 3D3, IgG1 | Abcam | ab140779 | Citrate | 200 |
| **iNOS** | M1 macrophages | Rabbit | Polyclonal | Abcam | ab3523 | Citrate | 400 |
| **CD163** | M2 macrophages | Mouse | EdHu-1, IgG1 | AbD-serotec | mca1853 | Citrate | 250 |
| **CD44** | Macrophage fusion | Rabbit | Polyclonal | Abcam | Ab24504 | Citrate | 250 |
| | Paracrine signaling | | | | | | |
| **TGF-β1** | Promotor of ECM | Rabbit | Polyclonal | Abcam | ab9758 | Citrate | 500 |
| **TNF-α** | Pro-inflammatory cytokine | Rabbit | Polyclonal | Acris | AP20372PU-n | Citrate | 300 |
| **IL-10** | Anti-inflammatory cytokine | Rabbit | Polyclonal | Biorbyt | orb221323 | Citrate | 200 |

#### Supplementary Figures

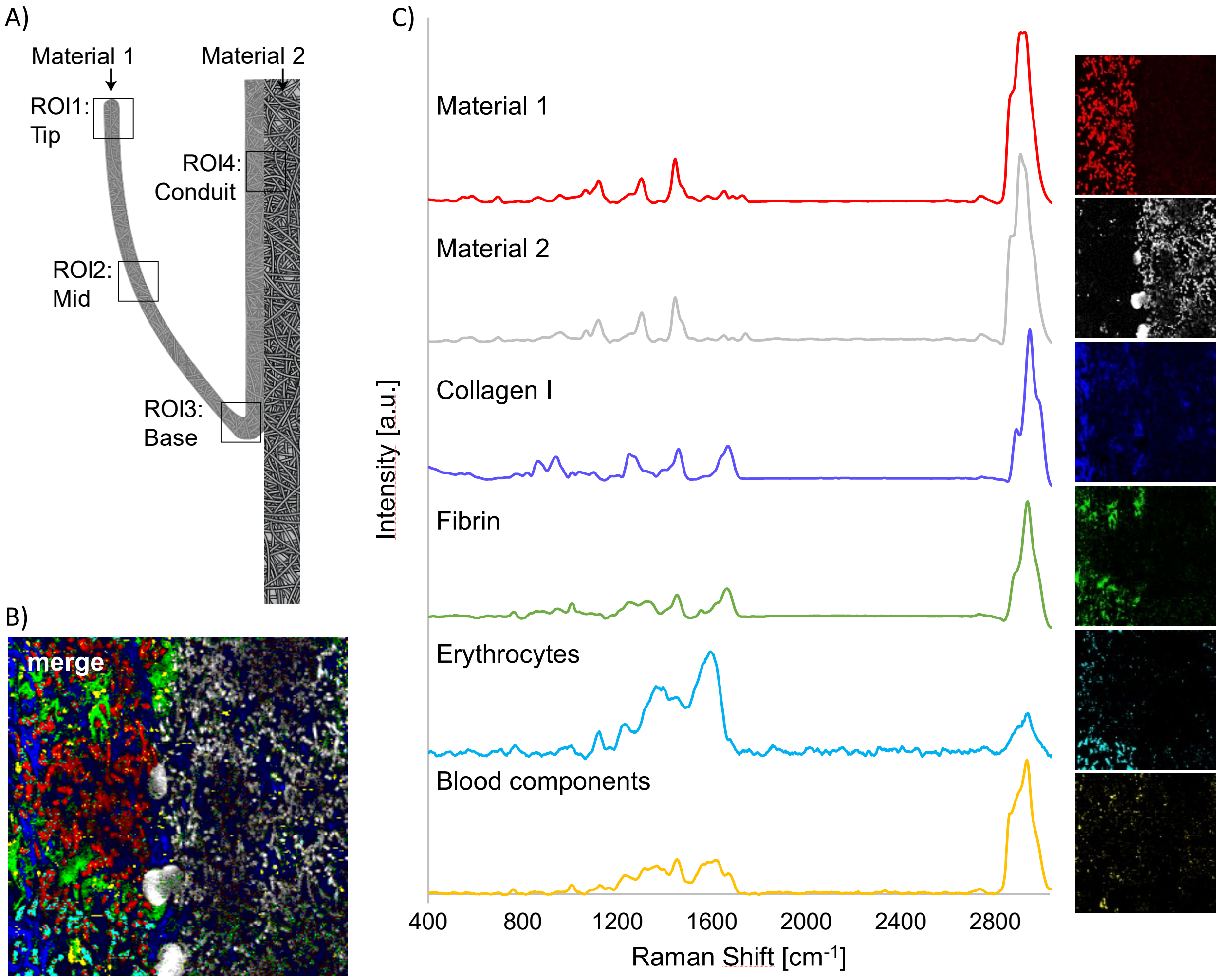


**Supplementary Figure S1:** A) Schematic overview of the graft with specified regions of interest (ROIs) used for Raman imaging; tip (ROI1), mid (ROI2) and base (ROI3) of the leaflet as well as the pulmonary conduit (ROI4). B) Example image of pseudo-color map of components as identified with True Component Analysis (TCA). C) TCA of the generated hyperspectral maps allowed for the identification and localization of the following components: implant material 1 (red) and material 2 (grey), collagen I (blue), fibrin (green), erythrocytes (cyan) and blood components (yellow).

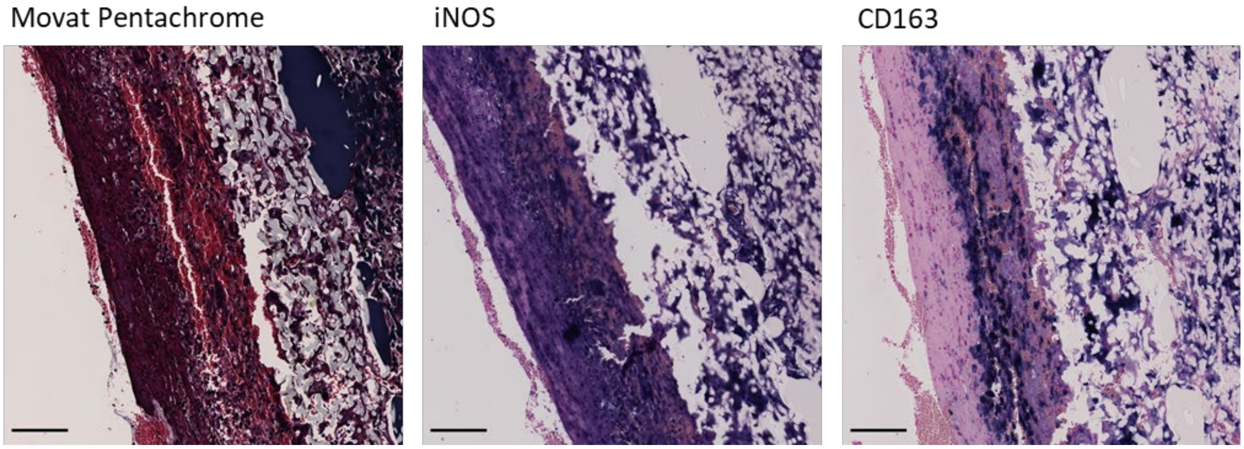


**Supplementary Figure S2: Erythrocyte-rich matrix on conduit of 2-month explant.** Images of Movat Pentachrome staining and immunohistochemistry staining for iNOS and CD163 of the neotissue on the conduit region as found in one explant (explant #CV36124) after 2 months of implantation. This region was rich in erythrocytes and rich in both anti- and pro-inflammatory macrophages. (Positive stain; dark purple, counterstain; pink). Scalebar equals 100 µm.

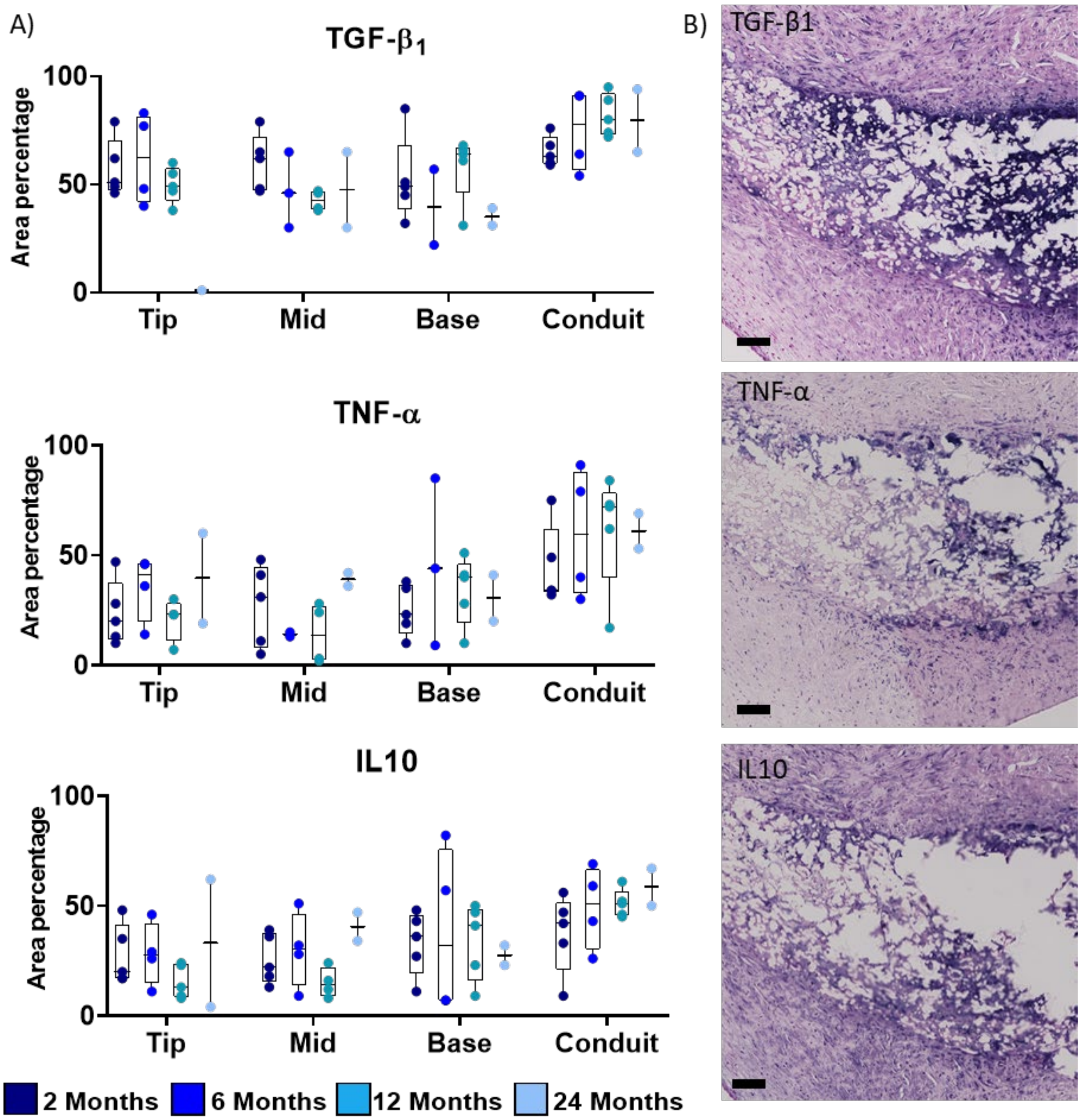


**Supplementary Figure S3: Analysis of cytokine secretion over time** A) Semi-quantification of immunohistochemical stainings (area 100.000 µm$^2$) of the transforming growth factor $\beta_1$ (TGF$\beta_1$), pro-inflammatory cytokine tumor necrosis factor α (TNFα) and anti-inflammatory cytokine interleukin 10 (IL10). B) Representative images of respective stainings of 6 month explants within base region. (Positive stain; dark purple, counterstain; pink). Scalebar equals 100 µm.

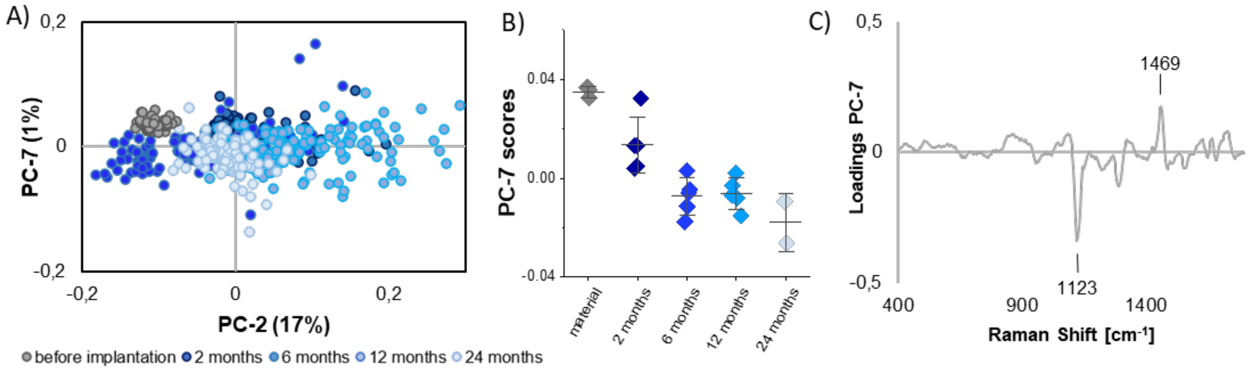


**Supplementary Figure S4: Raman analysis resorption material 2.** Principal Component Analysis (PCA) of the resorption of the conduit material 2 (ROI4). A) The PCA scores plot and B) the average score values show an implantation duration dependent shift in PC-7. C) Upon degradation, the loadings plot indicates an increase of the 1123 $cm^{-1}$ band and a decrease of the 1469 $cm^{-1}$ peak.

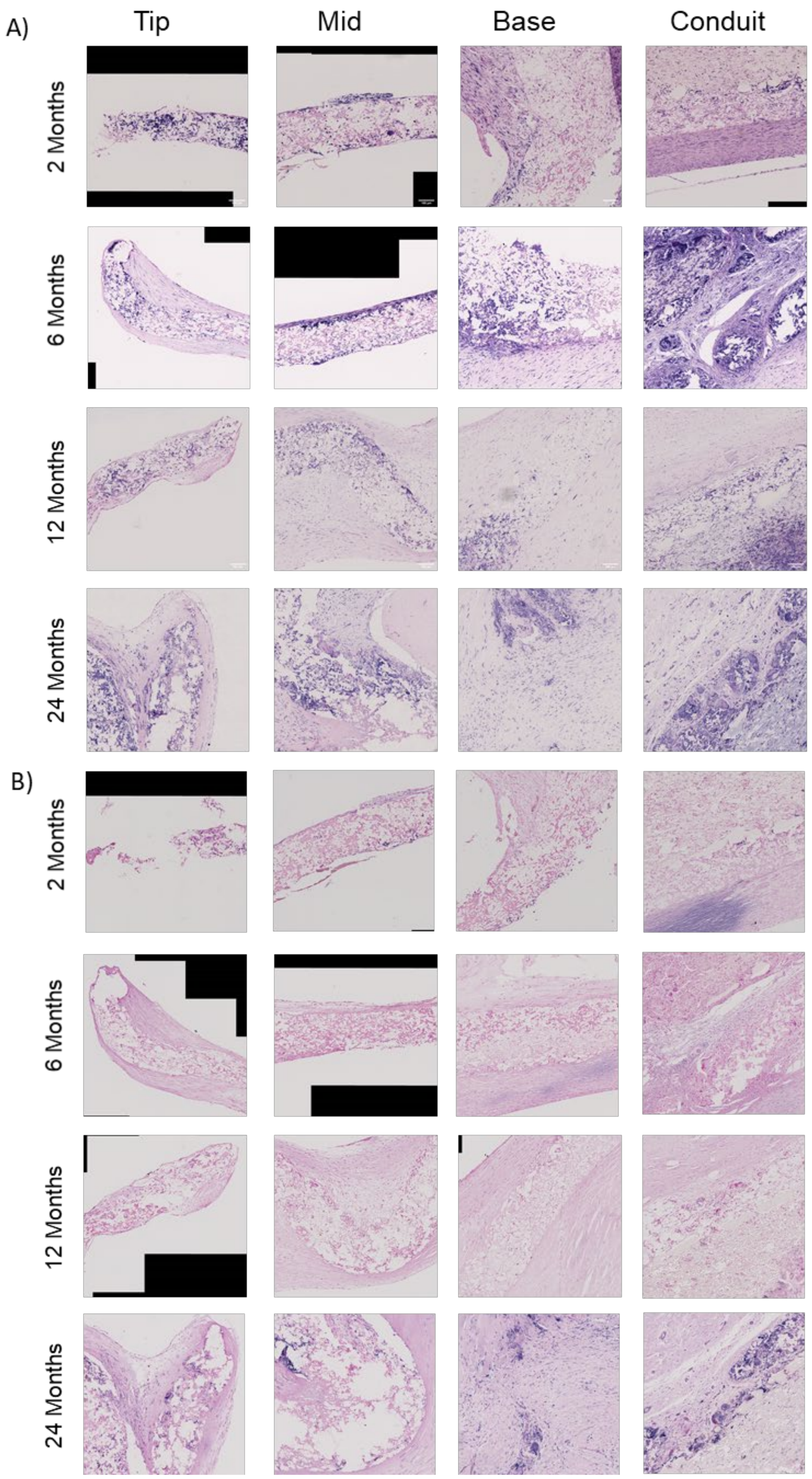


**Supplementary Figure S5: Immunohistochemical staining of CD44 and Vimentin.** Representative images of explants stained for CD44 (A) and Vimentin (B) for each timepoint and region of interest. (Positive stain; dark purple, counterstain; pink). Scalebar equals 100 µm.

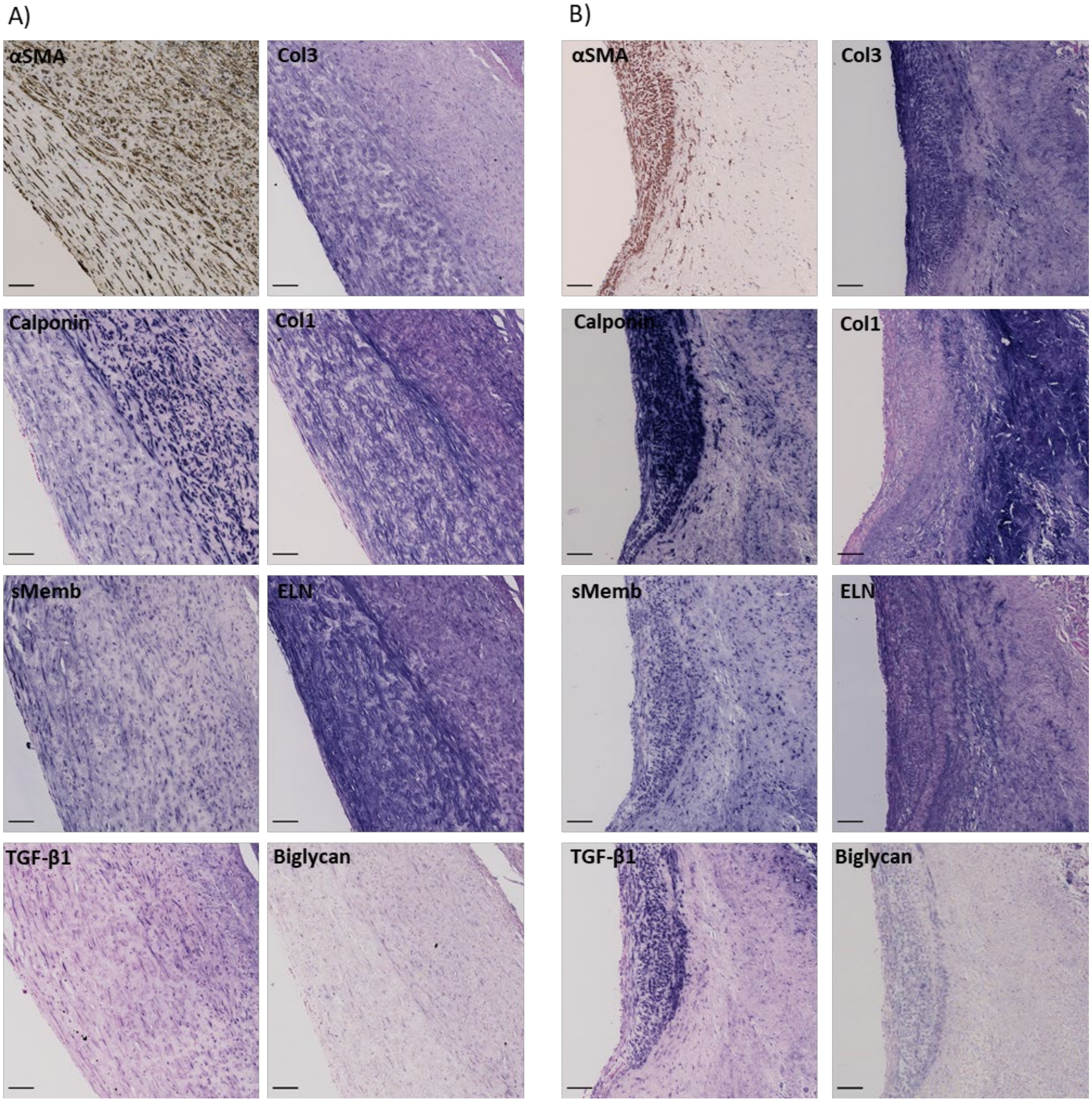


**Supplementary Figure S6: IHC of colocalizing tissue deposition of regions with specific VIC-like marker expression.** IHC images of αSMA, calponin and SMemb in the conduit region of a 6 month explant (A) and a 24 month explant (B) and IHC images of collagen 3 (Col3), collagen 1 (Col1), (tropo)elastin (ELN), biglycan and TGF-$\beta_1$ localizing within these respective regions. αSMA ; positive staining in brown, nuclei in blue, Other markers; positive staining in purple counterstain in pink. Scale bars, 100 µm.

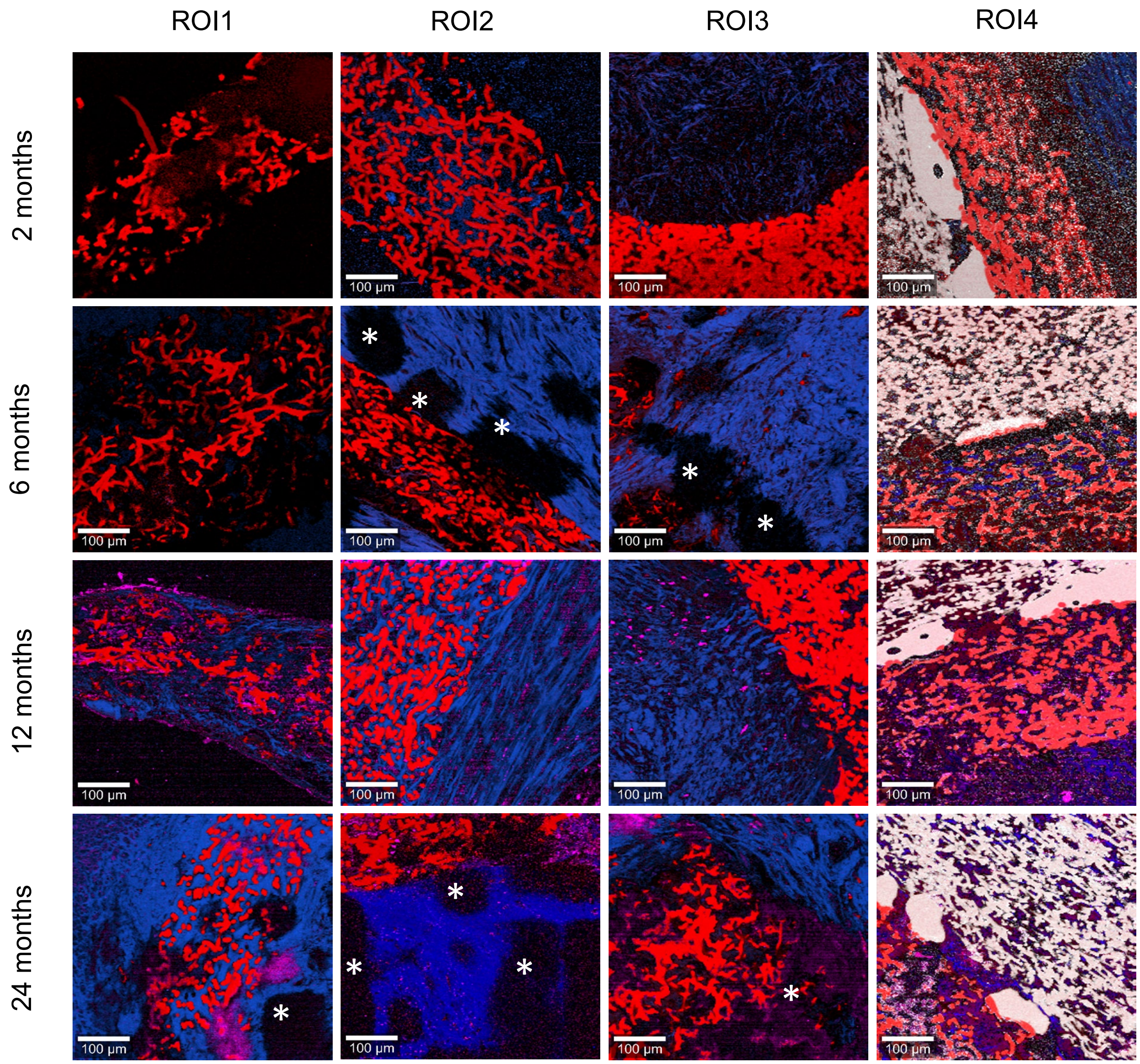


**Supplementary Figure S7: Overview of TCA images from Raman per ROI over time**. After 2 months little to no collagen was present within the graft material of the tip (ROI1) and mid (ROI2) regions of the leaflet. Within the base (ROI3) region of the leaflet and the graft material of the conduit (ROI4) collagen is present at all timepoints. Blue: collagen, red: material 1, white: material 2, magenta: nuclei. Scale bars equal 100 µm; asterisks indicate out-of-focus areas.

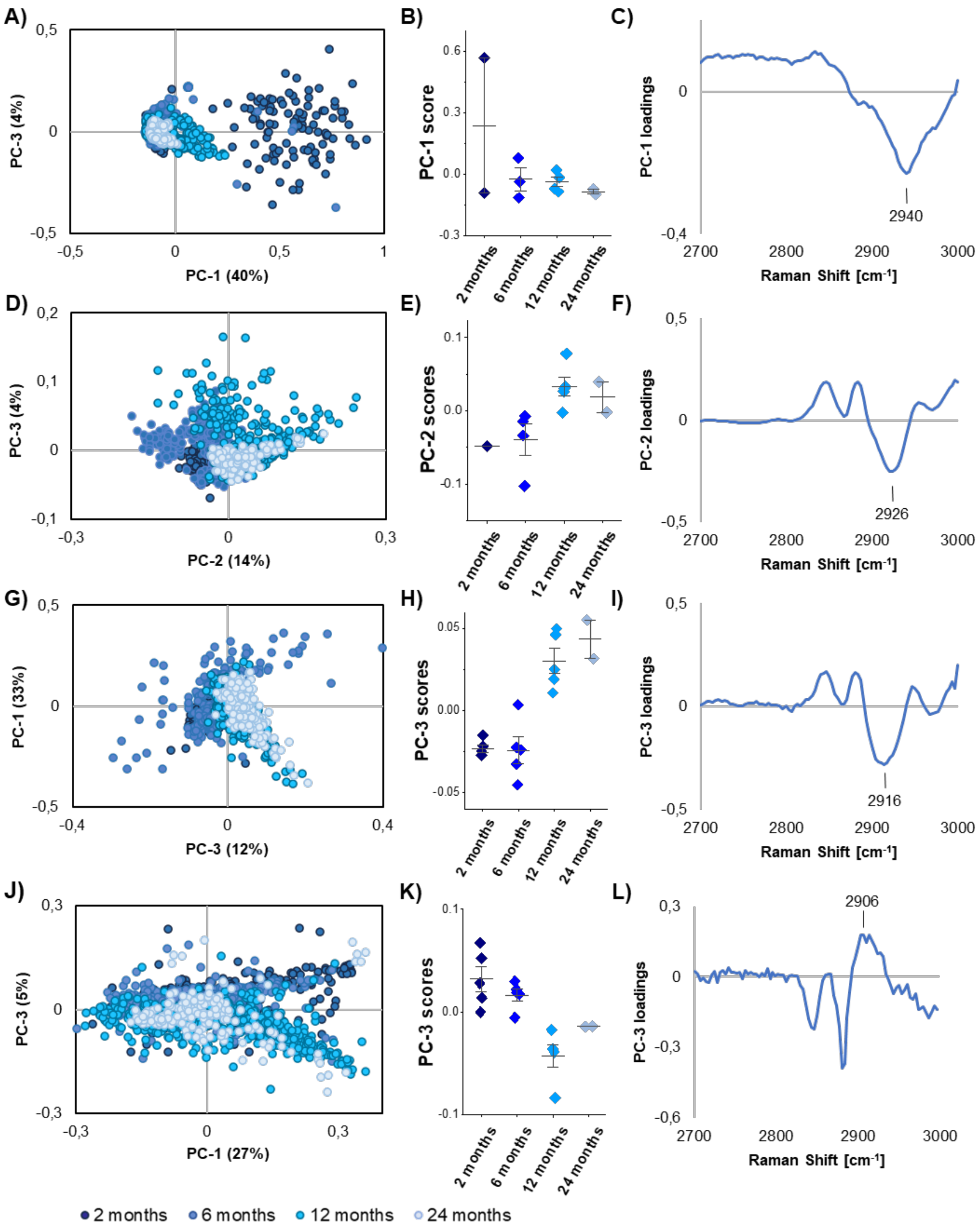


**Supplementary Figure S8: PCA Raman analysis for collagen maturation.** Raman analysis (high wavenumber region) of the collagen structures was performed to compare the different ROIs over time. (A-C) ROI1, (D-F) ROI2, (G-I) ROI3, and (J-L) ROI4. According to the loadings, differences in ROI1, ROI2 and ROI3 were mainly linked to the overall collagen content, whereas loadings for ROI4 were mainly linked to collagen maturation.

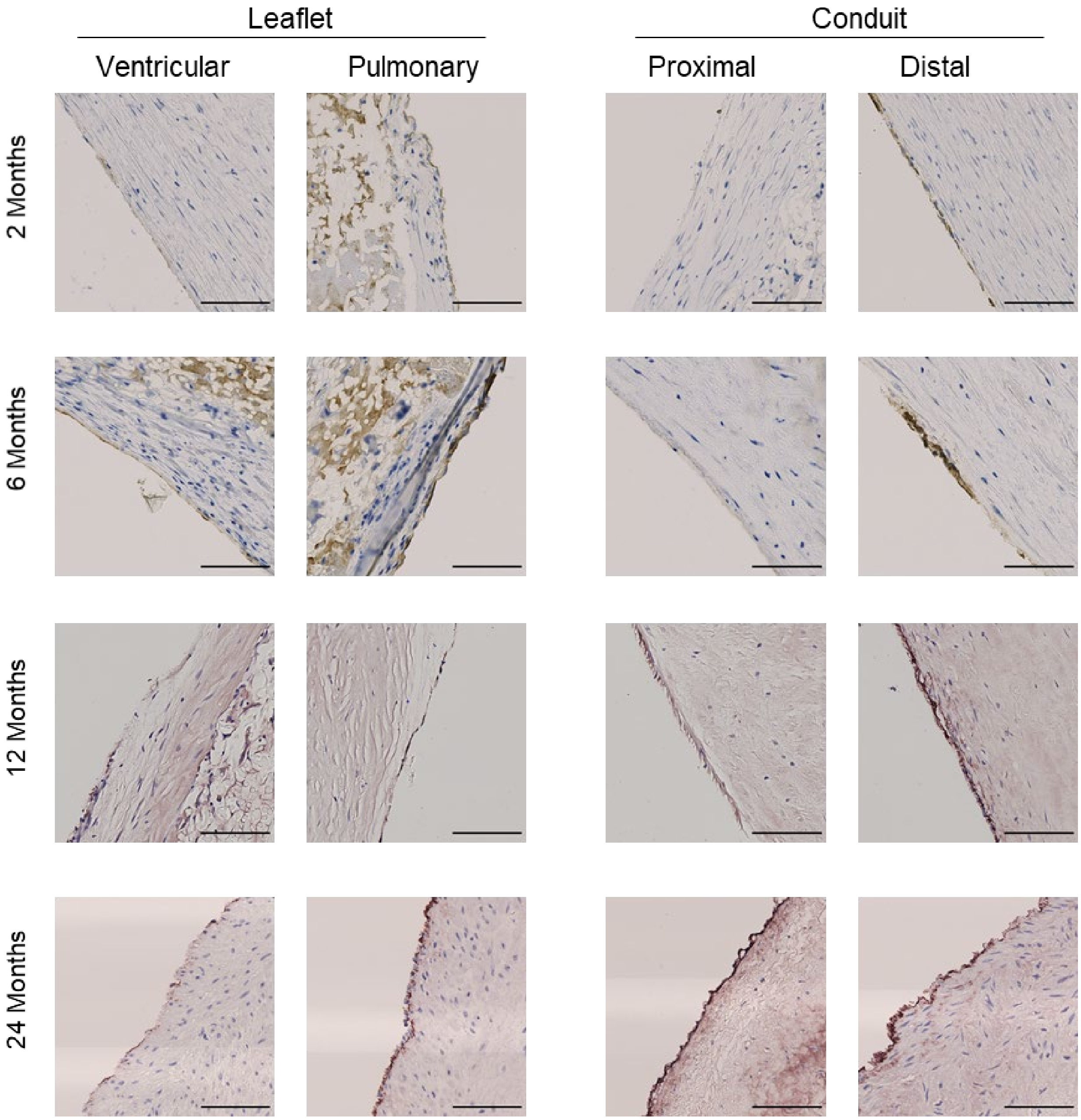


**Supplementary Figure S9: IHC of Endothelialization.** Representative images of the ventricular and pulmonary side of the base region of the leaflet and the proximal and distal side of the luminal surface of the neotissue deposited on the conduit after 2, 6, 12 and 24 months. Sectioned are stained for VonWillebrand Factor (vWF), indicative of endothelialization. After 2 months, vWF positive staining was present at the distal side of the conduit. Only sparse expression was present on the leaflet. After 12 months, in all regions the endothelial layer became more pronounced. Positive staining; brown, cell nuclei; blue. Scalebars equal 100 µm.

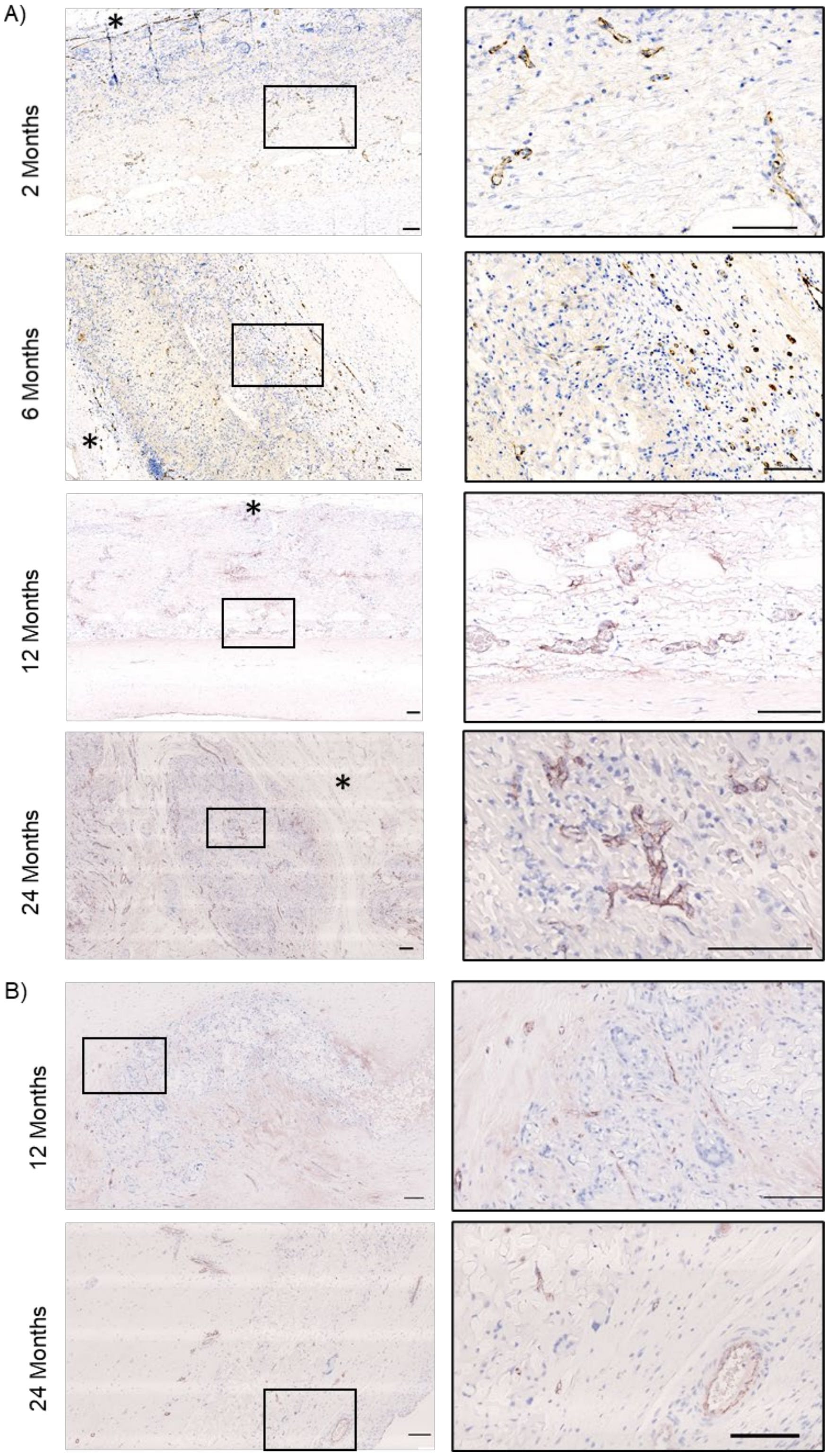


**Supplementary Figure S10: IHC of microvascularization.** Representative **i**mages and zooms of explants stained for VonWillebrand Factor indicating vascularization within the conduit material (A) after 2, 6, 12 and 24 months and vascularization in the leaflet (B) after 12 and 24 months. * indicates external conduit material. Positive staining; brown, cell nuclei; blue. Scalebars equal 100 µm.

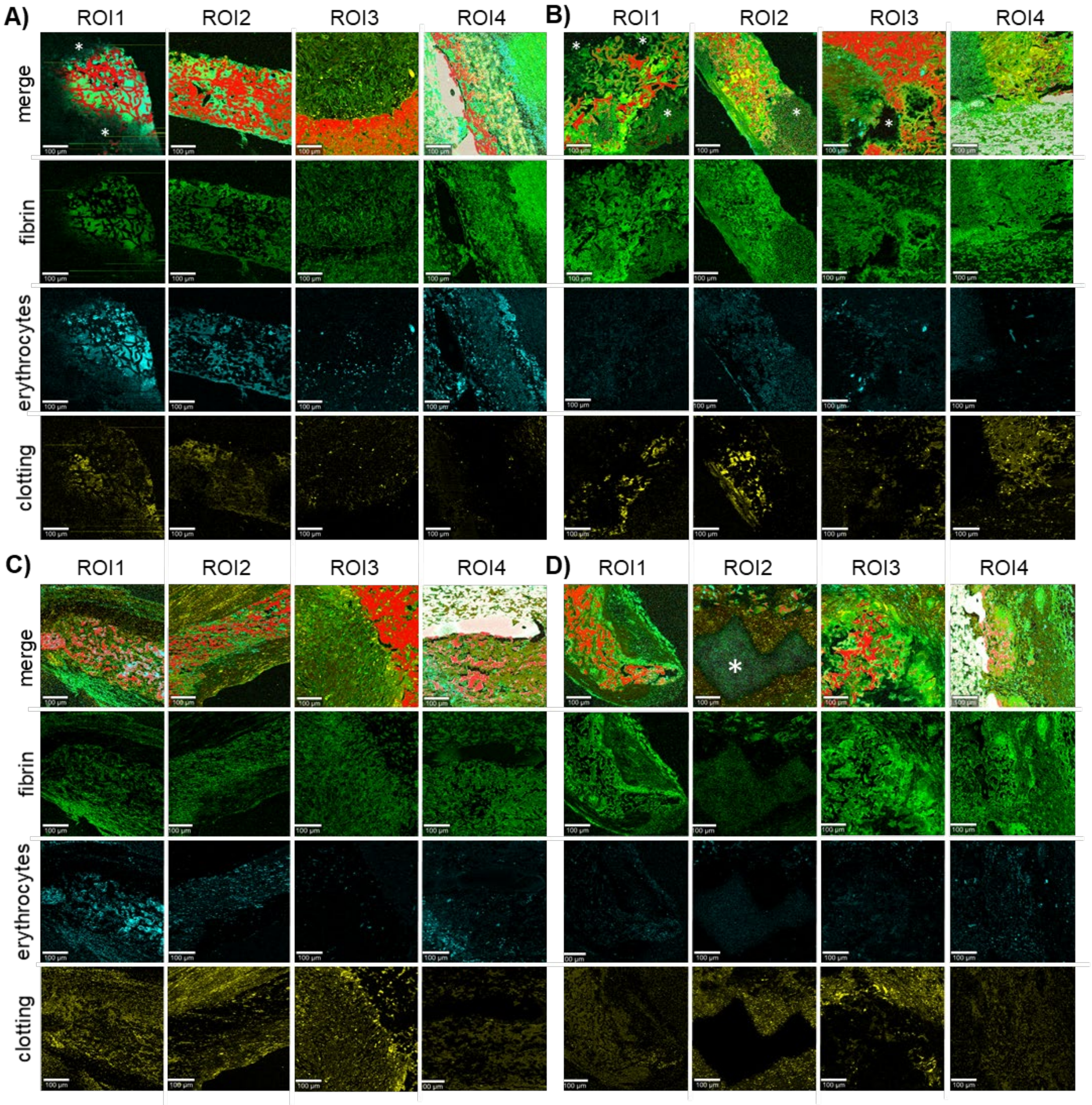


**Supplementary Figure S11: TCA Raman images of identified fibrin, erythrocytes and clotting material per ROI over time.** TCA identified the distribution of fibrin (green), erythrocytes (cyan) and blood components (yellow) in the different ROIs at (A) 2, (B) 6, (C) 12 and (D) 24 months. Red: material 1; White: material 2. Scale bars equal 100 µm; asterisks indicate out-of-focus areas.
